# Duplex-Repair enables highly accurate sequencing, despite DNA damage

**DOI:** 10.1101/2021.05.21.445162

**Authors:** Kan Xiong, Douglas Shea, Justin Rhoades, Tim Blewett, Ruolin Liu, Jin H. Bae, Erica Nguyen, G. Mike Makrigiorgos, Todd R. Golub, Viktor A. Adalsteinsson

**Affiliations:** Broad Institute of MIT and Harvard, Cambridge, MA, 02142, USA; Department of Radiation Oncology, Dana-Farber Cancer Institute and Brigham and Women’s Hospital, Harvard Medical School, Boston, MA; Department of Pediatric Oncology, Dana-Farber Cancer Institute, Boston, MA, USA

## Abstract

Accurate DNA sequencing is crucial in biomedicine. Underlying the most accurate methods is the assumption that a mutation is true if altered bases are present on both strands of the DNA duplex. We now show that this assumption can be wrong. We establish that current methods to prepare DNA for sequencing, via ‘End Repair/dA-Tailing,’ may substantially resynthesize strands, leading amplifiable lesions or alterations on one strand to become indiscernible from true mutations on both strands. Indeed, we discovered that 7-17% and 32-57% of interior ‘duplex base pairs’ from cell-free DNA and formalin-fixed tumor biopsies, respectively, could be resynthesized *in vitro* and potentially introduce false mutations. To address this, we present Duplex-Repair, and show that it limits interior duplex base pair resynthesis by 8- to 464-fold, rescues the impact of induced DNA damage, and affords up to 8.9-fold more accurate duplex sequencing. Our study uncovers a major Achilles’ heel in sequencing and offers a solution to restore high accuracy.

## INTRODUCTION

Mutations in DNA drive genetic diversity^1^, alter gene function^2^, impact cellular phenotypes^3^, mark cell populations^4^, define evolutionary trajectories^5^, underscore diseases and conditions^6^, and provide targets for precision medicines and diagnostics^7^. It is thus crucial to be able to detect mutations across a wide range of abundances. For instance, detecting low-abundance mutations (e.g. <0.1-1% VAF, down to ‘single duplex’ resolution) is important for studying cancer evolution^8^ and drug resistance^9^, understanding somatic mosaicism^10^ and clonal hematopoiesis^11^, characterizing base editing technologies^12^, evaluating the mutagenicity of chemical compounds^13^, uncovering pathogenic variants^14^, studying human embryonic development^15^, detecting microbial or viral infections^16^ and cancers^17^ and clinically actionable genomic alterations from specimens such as tissue or liquid biopsies^18^, and much more.

Despite progress in next generation sequencing (NGS), DNA damage confounds mutation detection and renders accuracy dependent upon sample quality, which is deeply problematic^19^. Lesions such as uracil, thymine dimers, pyrimidine dimers, 8-oxoGuanine (8’oxoG), 6-*O*-methylguanine, depurination, and depyrimidination arise both spontaneously and in response to environmental and chemical exposures, such as UV radiation, ionization radiation, reactive oxygen species, and genotoxic agents, or sample processing procedures, such as formalin fixation, freezing and thawing, heating and thermal cycling, acoustic shearing, and long-term storage in aqueous solution^20,21^. When amplified, translesion synthesis could occur, introducing a mutation *in vitro*. These, along with other errors in sample preparation and sequencing, contribute to an error rate of 0.1-1% in NGS^22^.

Due to the stochasticity of base damage errors, most can be overcome by barcoding and sequencing multiple copies of each DNA fragment and requiring a consensus among reads. Such methods can reduce errors by up to 100-fold, when requiring a consensus from each single strand of DNA, and up to 10,000-fold, when requiring a consensus from both sense strands of each DNA duplex in a technique called duplex sequencing^23^. However, most double-stranded DNA fragments, including those which have been sheared for sequencing, have ‘jagged ends’ which must be repaired in order to ligate sequencing adapters to both strands. ‘End Repair / dA-Tailing’ (ER/AT) methods are designed to remove 3’ overhangs, fill-in 5’ overhangs, phosphorylate 5’ ends (via ‘End Repair’), and leave a single dAMP on each 3’ end (via ‘dA-tailing’) to facilitate ligation of dTMP-tailed adapters. Yet, ER/AT methods include polymerases which may resynthesize portions of each duplex.

If resynthesis occurs in the presence of an amplifiable lesion or alteration confined to one strand, the altered base pairing will be propagated to the newly synthesized strands when amplified. This will render an amplifiable lesion or alteration from one strand indiscernible from a true mutation on both strands (**Fig. 1A**). This issue has been observed at the ends of each duplex (e.g. last ~12bp) due to fill-in of short 5’ overhangs^24^. However, we reason that such errors could also span much deeper given (i) the 5’ exonuclease and strand-displacement activities of Taq and Klenow polymerases used in ER/AT^25^ and (ii) the varied nicks, gaps, and overhangs in DNA^26^ which could act as ‘priming sites’ for strand resynthesis.

**Figure 1.**
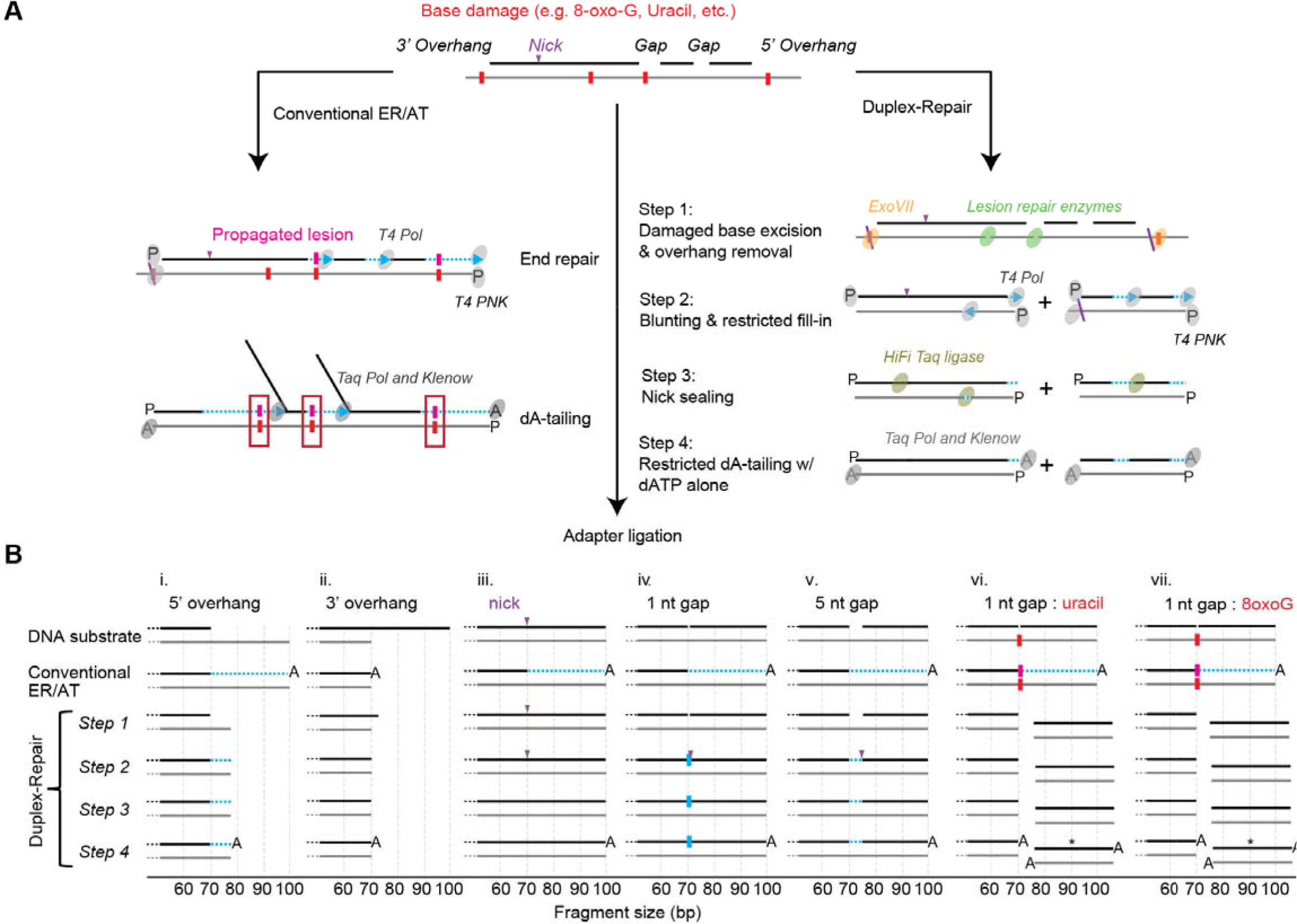
Characterization of Duplex-Repair using capillary electrophoresis. (A) overview of Duplex-Repair vs. conventional ER/AT methods (B) Schematic of the major products of various synthetic duplexes subjected to each step of Duplex-Repair and conventional ER/AT as determined by capillary electrophoresis (Raw traces are in Fig S2). The non-fluorophore-tagged ends of the synthetic molecules are depicted, and fragment sizes are drawn to scale. Duplexes demarcated by asterisks (*) do not contain fluorophores and were not directly observed by capillary electrophoresis; however, their presence is predicted due to the characterized activities of UDG and FPG. Regions of strand resynthesis are illustrated in light blue.

Here, we demonstrate that substantial portions of each duplex are resynthesized when conventional ER/AT is applied to DNA bearing nicks, gaps, or overhangs. We then describe a new ER/AT method called Duplex-Repair which limits strand resynthesis. Using single-molecule and panel sequencing, we show that Duplex-Repair minimizes strand resynthesis and restores high accuracy despite varied extents of DNA damage, when applied to samples such as cfDNA and formalin-fixed tumor biopsies.

## MATERIAL AND METHODS

### Duplex-Repair workflow

Duplex-Repair consists of four steps. In step 1, DNA is treated with an enzyme cocktail consisting of EndoIV (Cat. No. M0304S), Fpg (Cat. No. M0240S), UDG (Cat. No. M0280S), T4 PDG (Cat. No. M0308S), EndoVIII (Cat. No. M0299S) and ExoVII (Cat. No. M0379S) (all from NEB; use 0.2 uL each) in 1X NEBuffer 2 in the presence of 0.05 ug/uL BSA (total reaction volume = 20 uL) at 37 °C for 30 min. In step 2, T4 PNK (Cat. No. M0201S; NEB; use 0.25 uL), T4 DNA polymerase (Cat. No. M0203S; NEB; use 0.25 uL), ATP (final concentration = 0.8 mM), and dNTP mix (final concentration of each dNTP = 0.5 mM) are added into the step 1 reaction mix and incubated at 37 °C for another 30 min. In step 3, HiFi *Taq* ligase (Cat. No. M0647S; NEB; use 0.5 uL) and 10X HiFi *Taq* ligase buffer (use 1.5 uL) are spiked into the step 2 reaction mix and incubated on a thermal cycler that heats from 35 °C to 65 °C over the course of 45 min. The resulting products are purified by performing 3X Ampure bead cleanup and eluted in 17 uL of 10 mM Tris buffer. In step 4, the purified products are treated with Klenow fragment (3’ → 5’ exo-) (Cat. No. M0212L; NEB; use 1 uL) and *Taq* DNA polymerase (Cat. No. M0273S; NEB; use 0.2 uL) in 1X NEBuffer 2 in the presence of 0.2 mM dATP (total reaction volume = 20 uL) at room temperature for 30 min followed by 65 °C for 30 min. To prepare Duplex-Repair libraries for sequencing, T4 DNA ligase (Cat. No. M0202L; NEB; use 1000 units), 5’-deadenylase (Cat. No. M0331S; NEB; use 0.5 uL), PEG 8000 (final concentration = 10% (w/v)), and custom dual index duplex UMI adapters (IDT) are added to the step 4 reaction mix (total reaction volume = 55 uL) which is then incubated at room temperature for 1 hr followed by performing 1.2X Ampure bead cleanup, and the purified products are amplified by PCR.

### Quantification of strand resynthesis on synthetic oligonucleotides by capillary electrophoresis

fluorophore-labeled single-stranded oligonucleotides (from IDT; **Table S1**) were resuspended in low TE buffer (pH 8.0) and annealed to form DNA duplexes bearing nicks, gaps, or overhangs. Then, 20 - 200 ng of each duplex substrate was carried through the workflow of a conventional ER/AT kit, the Kapa Hyper Prep kit, or Duplex-Repair, and aliquots of products after each step were sent to Eton Bioscience for capillary electrophoresis analysis. The returned data were analyzed with Peak Scanner 2 software and then recalibrated (see **Supplementary text, Equation S1**, and **Equation S2**).

### Clinical specimens

All patients provided written informed consent to allow the collection of blood and/or tumor tissue and analysis of genetic data for research purposes. Healthy donor blood samples were ordered from Research Blood Components or Boston Biosciences. Patients with metastatic breast cancer were prospectively identified for enrollment into an IRB-approved tissue analysis and banking cohort (Dana-Farber Cancer Institute [DFCI] protocol identifier 05-055). Plasma was derived from 10-20cc whole blood in EDTA tubes.

### Quantification of strand resynthesis on cfDNA or gDNA by PacBio sequencer

We followed PacBio’s workflow for preparing multiplexed libraries by using the SMRTbell express template kit 2.0 (Pacific Biosciences) but made these modifications: 1). We skipped “Remove SS overhangs” and “DNA damage repair” steps; 2) We performed ER/AT by using the Kapa Hyper Prep kit or Duplex-Repair; 3). To perform ER/AT with d^6m^ATP (N6-methyl-2’-deoxyadenosine-5’-triphosphate), d^4m^CTP (N4-methyl-2’-deoxycytidine-5’-triphosphate), dGTP, and dTTP (all from TriLink Biotechnologies), we prepared and used a custom buffer (5x) consisting of 250 mM Tris, 2 mM d^6m^ATP, 2 mM d^4m^CTP, 2 mM dGTP, 2 mM dTTP, 50 mM MgCl_2_, 50 mM DTT, and 5 mM ATP (pH 7.5); 4). We performed 1.8X Ampure PB bead cleanup after nuclease treatment; 5). We skipped the “Second Ampure PB bead purification” step. The input into each library construction was 50 ng of a synthetic oligonucleotide or 20 - 40 ng of cfDNA or gDNA. As-prepared PacBio libraries were sequenced on Sequel II with a targeted read count of at least 65000 per sample.

### Induction of DNA damage by CuCl_2_/H_2_O_2_ and DNase I

We first optimized the conditions for inducing DNA damage by CuCl_2_/H_2_O_2_ and DNase I (Fig. S11-13 & Table S2). Then, 20 ng of cfDNA was treated with 0, 0.2, or 2 mU DNase 1 (Cat. No. M0303S; NEB) and 0, 1, or 100 uM CuCl_2_/H_2_O_2_ in 1X DNase 1 buffer (total reaction volume = 20 uL) at 16 °C for 1 hr. 40 mM EDTA was then added to quench the reaction, and the resulting products were purified by performing a 2X Ampure bead cleanup.

### Processing of cfDNA sample and gDNA sample

cfDNA was extracted from fresh or archival plasma of healthy donors or cancer patients by following the same method as before^24,27^. gDNA was extracted from FFPE tumor tissues or buffy coats, sheared and quantified by following the same protocol as previously described^24,27^. Then, cfDNA or gDNA libraries were constructed from 10-20 ng DNA inputs by using the Kapa Hyper Prep kit or Duplex-Repair with custom dual index duplex UMI adapters (IDT). Hybrid Selection (HS) using IDT’s pan-cancer panel was performed on the prepared libraries using the xGen hybridization and wash kit with xGen Universal blockers (IDT). After the second round of HS, libraries were amplified, quantified and pooled for sequencing on a HiSeq 2500 rapid run (100 bp paired-end runs) or HiSeqX (151 bp paired-end runs) with a targeted raw depth of 200,000x per site.

### Analysis of duplex sequencing data and quantification of error rates

Raw reads were then processed through our duplex consensus calling pipeline as previously described^24^. We calculated error rates by counting the proportion of non-reference bases to total bases after applying filters specifically tailored to duplex sequencing^24^. To avoid miscounting true somatic variants from cancer patients as base errors, we omitted any loci that had a somatic mutation called from whole exome sequencing of that patient’s tumor biopsy. We also used a matched normal derived from buffy coat DNA to filter any germline mutations. For base error position analysis, we reran our error metrics collection pipeline with the end of fragment filter disabled to observe errors across the entire DNA duplex.

### Estimating resynthesis from single molecule real-time (SMRT) sequencing data

We first used the Circular Consensus Sequences (CCS) tool (Pacific Biosciences) to generate consensus reads from the raw reads. We also used the --mean-kinetics flag to output interpulse durations (IPDs), among other metrics, for each base position to be used later for identifying modified dNTPs. We then used the lima tool (Pacific Biosciences) to demultiplex the samples that were sequenced together on the same flow cell. These CCS reads were then used as input for our Hidden Markov Model (HMM) to estimate strand resynthesis.

We implemented a HMM to estimate the amount of resynthesis on the 3’ end of each duplex strand from SMRT sequencing data. The HMM consists of two states that represent regions with original bases (O) and regions with bases that were filled-in during ER/AT (F) respectively. We designed the HMM to estimate resynthesis that starts at an interior position in the strand and continues all the way to the 3’ end. In addition, we designed a transition matrix that does not allow F to O transitions. We then set the transition probability from O to F, x, equal to the reciprocal of the strand length and the transition probability from O to O, y equal 1-x. To develop an empirical emission matrix, we sequenced synthetic duplexes with known regions of resynthesis and of original bases(**Table S1**). PacBio SMRT sequencing emits both the base and interpulse duration (IPD) for each position which we then collected to form the emission matrix of IPD distributions for each base in each state (**Fig. S12**). Using this HMM, we applied the Viterbi algorithm to each duplex DNA strand to determine the most likely regions of original bases and of resynthesized bases and calculated the total number of resynthesized bases.

To estimate the fraction of interior base pair resynthesized, we took the regions of estimated resynthesis from our HMM and counted the number of resynthesized base pairs that were greater than 12 base pairs from either end of the duplex fragment relative to the number of total base pairs that were greater than 12 base pairs from either fragment end. For all analyses we also ran control samples with standard, non-modified dNTPs to measure the background resynthesis estimates and subtracted that background from our samples where modified dNTPs were used.

## RESULTS

### Duplex-Repair as a new ER/AT approach

We first wanted to test our hypothesis that conventional ER/AT methods could resynthesize substantial portions of DNA duplexes bearing nicks, gaps, or overhangs, including those with amplifiable lesions. To do so, we generated duplex oligonucleotides bearing (i) 5’ overhangs, (ii) 3’ overhangs, (iii) nicks, (iv-v) gaps of varied lengths without base damage, or (vi-vii) gaps with base damage (**Fig. 1B, Table S1**). The top and bottom strands were labeled with different dyes so that we could use capillary electrophoresis to quantify changes in fragment length during ER/AT (**Supplementary text; Fig. S1**). We applied conventional ER/AT methods and observed substantial strand resynthesis in all substrates except for those with 3’ overhangs (**Fig. 1B, Fig S2**). For instance, with even just a single nick in the middle of the top strand, the 30 bases downstream of the nick site were entirely resynthesized. Our results confirm conventional ER/AT methods can resynthesize large portions of each duplex, when nicks, gaps, or overhangs are present.

To address this issue, we devised a new approach called Duplex-Repair, which conducts ER/AT in a careful and stepwise manner to limit strand resynthesis (**Fig. 1A**). Duplex-Repair was designed to ‘concentrate’ resynthesis at fragment ends (e.g. last 12 bp) where errors can be trimmed *in silico^24^*. Duplex-Repair consists of four steps: (1) damaged base excision and overhang removal, (2) blunting and restricted fill-in, (3) nick sealing, and (4) restricted dA-tailing. In step 1, DNA is treated with an enzyme cocktail consisting of enzymes involved in Base Excision Repair (BER), such as Endonuclease IV (EndoIV), Formamidopyrimidine [fapy]-DNA glycosylase (Fpg), Uracil-DNA glycosylase (UDG), T4 pyrimidine DNA glycosylase (T4 PDG), and Endonuclease VIII (EndoVIII). These enzymes excise damaged bases such as Uracil, 8’oxoG, oxidized pyrimidines, cyclobutane pyrimidine dimers and cleave abasic sites, resulting in 1 nt gaps in double-stranded regions or strand breaks in single-strand regions. Exonuclease VII (ExoVII) is also used in this step to degrade 3’ and 5’ single-strand overhangs. Then, in step 2, T4 polynucleotide kinase (de)phosphorylates DNA termini, while T4 DNA polymerase blunts 3’ overhangs and fills in the small gaps and short (≤ 7 nt) 5’ overhangs which remain after ExoVII digestion. After that, nicks are sealed by HiFi *Taq* DNA ligase in step 3. In step 4, restricted dA-tailing is performed using Klenow fragment (exo-) and *Taq* DNA polymerase, but with only dATP present, to limit their activities to non-templated extension.

Using the aforementioned synthetic duplexes, we confirmed that Duplex-Repair facilitates ER/AT with minimal resynthesis. We first tested each step in ideal buffer conditions by performing a 3X Ampure bead cleanup after each step and have depicted the major products (**Fig. 1B & S2**). For each substrate, we confirmed the activity of the key enzymes involved, while making sure that the other enzymes present did not compromise their activity. For instance, for substrate (i), the long 5’ overhang is largely digested by ExoVII (**Fig. S3**) while the remaining three bases are filled in by T4 DNA polymerase (**Fig. S2**). For substrate (ii), the 3’ overhang is digested in part by ExoVII (**Fig. S3**), and then blunted entirely by T4 DNA polymerase (**Fig. S2**). For substrate (iii), the nick is sealed by HiFi Taq DNA ligase (**Fig. S4**). For substrates (iv-v), the gaps are first filled by T4 DNA polymerase (**Fig. S2 & S5**) and then the resulting ‘nicks’ are sealed by HiFi Taq DNA ligase. For substrates (vi-vii), the damaged bases are excised (uracil by UDG; 8’oxoG by Fpg; **Fig. S2 & S3**) and abasic sites cleaved to create strand breaks and thus avoid translesion synthesis during gap filling in step 2. We also confirmed that dA-tailing works with only dATP present (**Fig. S6 - S8**). We then optimized the reaction conditions and eliminated multiple Ampure cleanups between steps that would help reduce DNA loss (**Fig. S9 & S10**). Our results suggest that Duplex-Repair conducts ER/AT in a manner which limits strand resynthesis while achieving comparable library conversion efficiencies as conventional ER/AT (**Fig. S11**).

### Duplex-Repair limits resynthesis of DNA duplexes from clinical specimens

We next sought to quantify strand resynthesis when ER/AT is applied to clinical samples such as cell-free DNA (cfDNA) and formalin-fixed paraffin-embedded (FFPE) tumor biopsies. We devised an assay which involved performing ER/AT using a modified dNTP mix comprising d^6m^ATP, d^4m^CTP, dTTP, and dGTP, sequencing the prepared libraries on a PacBio sequencer which can detect where d^6m^ATP and d^4m^CTP have been incorporated^28^, and applying a Hidden Markov Model to identify resynthesized regions (**Fig. 2A & Fig. S12; Methods**). We first verified its performance using synthetic oligonucleotides (**Table S1**) treated with conventional ER/AT. We observed extended interpulse durations (IPDs) corresponding to d^6m^ATP and d^4m^CTP incorporations in the anticipated regions (**Fig. S13-i**). We also found the estimated number of resynthesized bases to be expected in most cases (**Fig. S13-ii**). Interestingly, for the substrates with a nick or a gap, we found some molecules with longer than expected fill-in, despite having the same terminal 3’OH as the substrate with a 80 bp 5’ overhang. We reason that this could be due to 3’ exonuclease activity of the polymerase, which may be pronounced when it encounters an adjacent, downstream strand.

**Figure 2:**
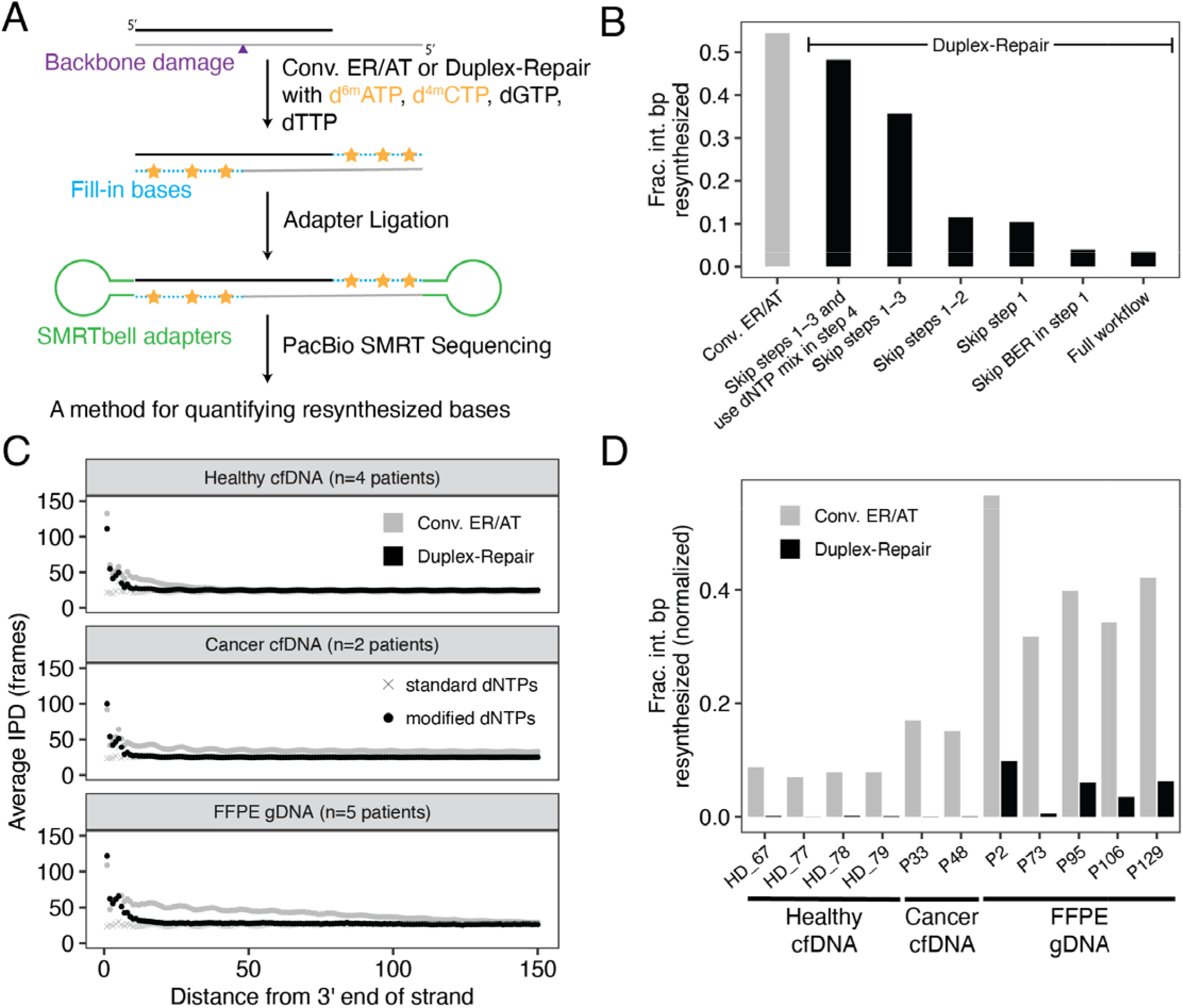
Quantification of strand resynthesis using Single-Molecule Real-Time (SMRT) sequencing. (A) Schematic of library construction for PacBio SMRT sequencing using modified dNTPs to aid in identifying resynthesis regions. (B) Estimated fractions of interior base pairs (> 12 bp from either end of the original duplex fragment) that were resynthesized using conventional ER/AT and several variations of Duplex-Repair. (C) Observed average interpulse durations (IPD; in frames) for circular consensus sequence (CCS) read strands relative to the distance from the original 3’ end of those strands across three sample types. (D) Estimated fraction of interior base pairs resynthesized for both conventional ER/AT and Duplex-Repair across three sample types.

We then used the above resynthesis quantification method to estimate the difference in resynthesized base pairs between Duplex-Repair and conventional ER/AT by testing on a healthy donor cfDNA sample with base and backbone damage induced by 100 uM CuCl_2_/H_2_O_2_ and 2 mU DNase 1, respectively (see **Methods**). We also tested several variations of Duplex-Repair in order to assess the impact of each step on limiting resynthesis. Applying our method, we estimated that 54% of interior duplex base pairs (defined as base pairs that are greater than 12 base pairs from either end of the original duplex DNA fragment) were resynthesized with conventional ER/AT, as compared to 3% with Duplex-Repair (**Fig. 2B**). Notably, each step in the Duplex-Repair protocol we tested served to reduce the amount of interior base pair resynthesis further. In particular, we observed that skipping the BER in step 1 had a negligible impact on resynthesis while skipping step 1 increased interior resynthesis fractions from 3% to 9%, suggesting that ExoVII treatment is required for suppressing resynthesis on 5’ overhangs. Further, skipping step 2 only slightly increased interior resynthesis fractions from 9% to 11%, confirming limited resynthesis occured during restricted fill-in. Further, skipping step 3 increased interior resynthesis fraction from 11% to 35%, suggesting that unsealed nicks led to significant resynthesis during dA-tailing. Furthermore, using dNTP mix instead of dATP alone in step 4 increased the resynthesis fraction from 35% to 47%, suggesting that it is essential to use dATP alone to suppress templated extension during dA-tailing. Overall, these results suggest that the full protocol of our Duplex-Repair is required to minimize resynthesis.

To assess the extent to which Duplex-Repair could limit resynthesis in clinical samples, we then used our assay to measure resynthesis across several different sample types, including healthy donor cfDNA, cancer patient cfDNA, and tumor FFPE biopsies. Considering that d^6m^ATP and d^4m^CTP could be present as real epigenetic modifications in clinical samples^29^, we also ran a control sample for each patient using all standard dNTPs and conventional ER/AT to control for any background noise. We first looked at average IPDs across strand positions for each CCS strand relative to the distance from the 3’ end of the original DNA strand (**Fig. 2C, Fig. S14**). For all sample types, we observed consistently low average IPDs across all positions for control samples. In contrast, average IPDs significantly increased both for conventional ER/AT and Duplex-Repair towards the 3’ ends of CCS strands (**Fig. 2C**). Furthermore, elevated IPDs for Duplex-Repair are concentrated within 12 base pairs from the 3’ end, but they extend much further into the strand for conventional ER/AT. Next we used our resynthesis quantification method to estimate the amount of interior duplex base pair resynthesis in our clinical samples. The fractions of interior base pairs resynthesized (after subtracting out the background noise from our control samples; **Fig. S15**) are much higher for conventional ER/AT compared to Duplex-Repair across all sample types (**Fig. 2D**). In particular, we observed that with conventional ER/AT, on average 8% (range 7-9%), 16% (range 15-17%), and 41% (range 32-57%) of interior duplex base pair resynthesis occurred for healthy cfDNA, cancer patient cfDNA, and FFPE tumor gDNA samples, respectively, which decreased to 0.12% (range 0.00-0.17%), 0.0345% (range 0.03-0.04%), and 5% (range 0.5-10%) when Duplex-Repair was used and thus corresponded to reductions in interior base pair resynthesis of 67-fold, 464-fold, and 8-fold. Our results suggest that conventional ER/AT induces substantial strand resynthesis in clinical samples such as cfDNA and FFPE tumor biopsies and that Duplex-Repair can significantly limit this.

### Duplex-Repair overcomes induced DNA damage and enhances duplex sequencing

Reasoning that strand resynthesis in ER/AT would be most problematic when amplifiable lesions or alterations are present, we subjected cfDNA from one healthy donor (HD_78) to different concentrations of the oxidizing agent CuCl_2_/H_2_O_2_, and DNase I to induce base and backbone damage without appreciably degrading DNA (**Fig. S16-S18 & Table S2**). We then applied conventional ER/AT, performed duplex sequencing, and computed error rates after trimming the last 12bp from the ends of each duplex^24^ (**Fig. 3A**, **Fig S19, Table S4**). At each concentration of CuCl_2_/H_2_O_2_, we found that the error rate increased with increasing amounts of DNase I, while the highest concentrations of both yielded an error rate 3.6-fold higher (C.I. 2.8-4.5) than that of untreated cfDNA. Expectedly, we observed the largest increase in errors which matched the expected C->A mutation signature of CuCl_2_/H_2_O_2_ exposure (13.9-fold, **Fig. S19**)^30^. Our results suggest that with conventional ER/AT, sequencing accuracy depends upon the extent of DNA damage in a sample.

**Figure 3.**
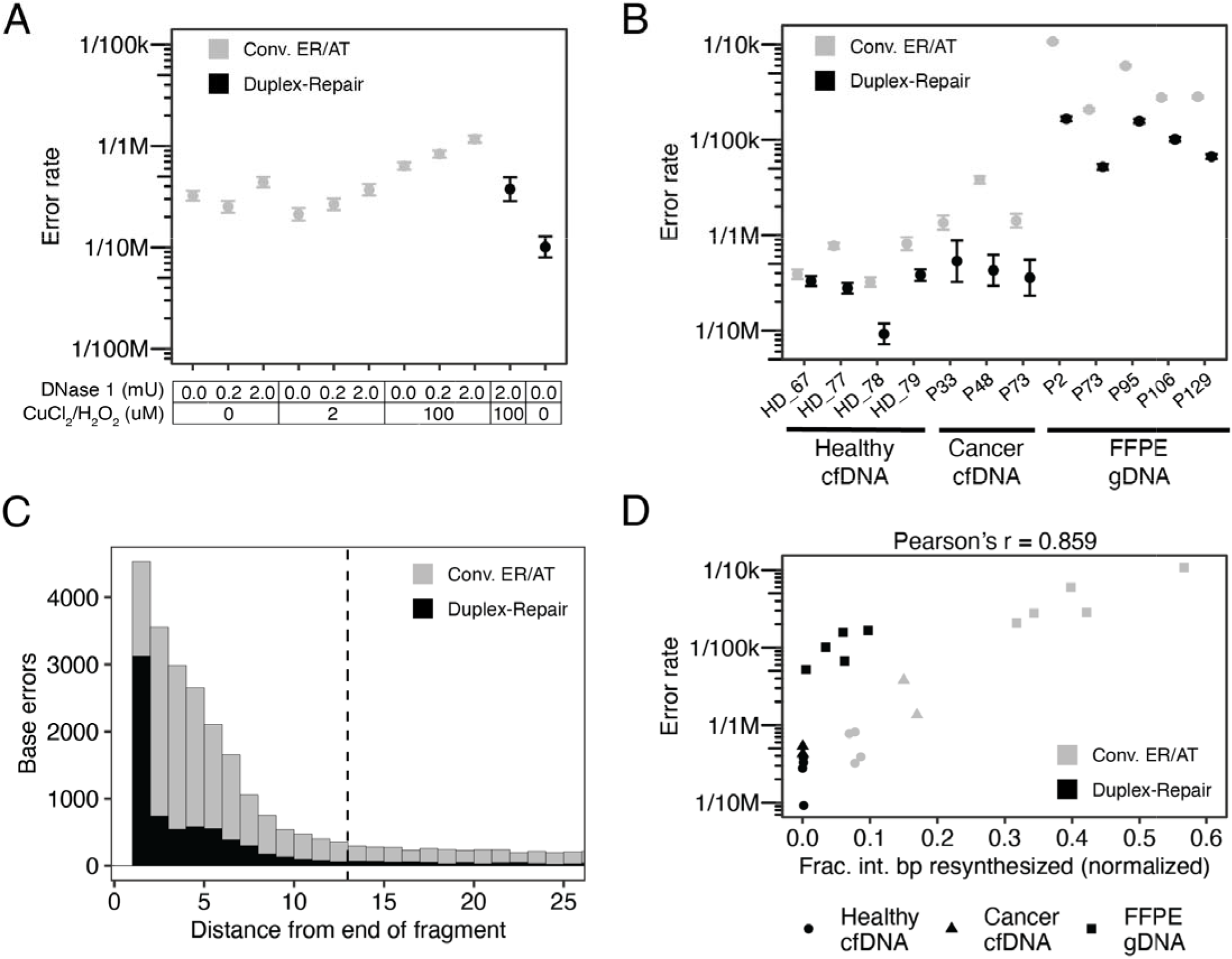
Targeted panel sequencing of cfDNA and FFPE tumor biopsies. (A) Measured duplex sequencing error rates of HD_78 cfDNA damaged with varied concentrations of DNase I (to induce nicks) and CuCl_2_/H_2_O_2_ (to induce oxidative damage) and then repaired by using Duplex-Repair or conventional ER/AT (three replicates per condition). (B) Duplex sequencing error rates of four healthy cfDNA samples (three replicates per condition), three cancer patient cfDNA samples (one replicate per condition), and five cancer patient FFPE tumor biopsies (three replicates per condition) treated with conventional ER/AT or Duplex-Repair. (C) Aggregate mutant bases and their position relative to the end of the original duplex fragment. Dashed line represents the threshold of the interior of the fragment (12bp). (D) Error rates from (B) compared to their corresponding estimates of interior base pair resynthesis fractions from Fig. 2D. Pearson’s correlation calculated for all data points.

To determine whether we could revert the impact of induced damage, we applied Duplex-Repair to the most heavily damaged samples and sequenced them with the same gene panel. We observed a significant reduction in error rate from 1.2e-6 to 3.7e-7, which was similar to the native cfDNA samples treated with conventional ER/AT (3.2e-7, **Fig. 3A**). Indeed, the impact of induced C->A errors was almost entirely ‘rescued’ (**Fig. S19**), while there was little change in error rates for other contexts (**Fig. S19**). We then applied Duplex-Repair to the native (i.e. undamaged) cfDNA and found the lowest error rates of all conditions tested (1.0e-7, **Fig. 3A**, **Fig. S19**). Our results suggest that Duplex-Repair can revert the impact of induced DNA damage.

Then, we sought to determine whether Duplex-Repair could provide higher accuracy than conventional ER/AT when used for duplex sequencing of clinical samples. We applied a 127-gene “pan-cancer” panel across three sample types (**Fig. 3B**). In all samples, we observed lower error rates when Duplex-Repair was applied, in comparison to conventional ER/AT. In particular, the median error rates decreased from 5.8e-7 (range 3.2e-7 - 8.1e-7) to 3.0e-7 (range 9.2e-8 - 3.8e-7) for healthy cfDNA, from 1.4e-6 (range 1.4e-6 - 3.8e-6) to 4.3e-7 (range 3.6e-7 - 5.3e-7) for cancer cfDNA, and from 2.8e-5 (range 2.1e-5 - 1.1e-4) to 1.0e-5 (range 5.2e-6 - 1.7e-5) for FFPE tumor biopsies, which amounts to a median 2.5-fold (C.I. 1.6 - 3.3), 4.0-fold (C.I. 3.4 - 4.5), and 4.0-fold (C.I. 3.1 - 4.9) reduction in error rates respectively, with cancer patient cfDNA from P48 showing the largest 8.9-fold reduction in error rate (**Fig. 3B**). Furthermore, the most significant reductions in duplex sequencing error rates occurred for contexts of C->T (median 3.6-fold, C.I. 2.5 - 4.1 for healthy cfDNA; median 5.7-fold, C.I. 5.3 - 5.8 for cancer cfDNA; median 4.1-fold, C.I. 3.1 - 5.0 for FFPE biopsies), C->A (median 3.4-fold, C.I. 2.7 - 3.8 for healthy cfDNA; median 3.8-fold, C.I. 3.6 - 4.0 for cancer cfDNA; median 19.0-fold, C.I. 18.7 - 19.3 for FFPE biopsies), and C->G (median 1.9-fold, C.I. 1.2 - 2.5 for healthy cfDNA; median 1.5-fold, C.I. 1.0 - 1.9 for cancer cfDNA; median 6.2-fold, C.I. 5.8 - 6.6 for FFPE biopsies; **Fig. S20, Table S3**). Notably, we observed that base errors were more significantly enriched at the ends of fragments with 34% of a total of 9,122 base errors (after normalizing for total bases evaluated) being in the first base from either duplex fragment end for Duplex-Repair as compared to only 15% of a total of 31,100 base errors for conventional ER/AT (**Fig. 3C, Fig. S21**). Overall, we estimated that 74% of base errors were concentrated within 12 bp from the end of the fragment for Duplex-Repair, in contrast with 68% for conventional ER/AT. It is worth noting that these base errors can be removed in-silico by filtering regions less than 12bp from the duplex fragment ends. Finally, we examined the relationship between strand resynthesis fractions and observed error rates across our clinical samples. We observed a strong overall correlation between the fractions of interior base pairs resynthesized and the error rates of duplex sequencing (Pearson’s r = 0.859; **Fig. 3D**). Our results establish that Duplex-Repair could afford consistently higher accuracy for duplex sequencing of clinical samples by limiting resynthesis during library construction.

## DISCUSSION

We have shown that existing ‘End Repair/dA-tailing’ (ER/AT) methods could resynthesize large portions of each DNA duplex, particularly when there are interior nicks, gaps, or long 5’ overhangs. This is a major problem for techniques such as duplex sequencing which require a consensus of reads from both strands. We then present a solution called Duplex-Repair which conducts ER/AT in a careful, stepwise manner. We show that it limits resynthesis by 8- to 464-fold, reverts the impact of induced DNA damage, and confers up to 8.9-fold higher accuracy in duplex sequencing of a cancer gene panel for specimens such as cfDNA and FFPE tumor biopsies. Considering the widespread use of duplex sequencing in biomedical research and diagnostic testing, our findings are likely to have broad impact in many areas such as oncology, infectious diseases, immunology, prenatal medicine, forensics, genetic engineering, and beyond.

Our study has characterized this major Achilles’ heel in ER/AT and provided a solution to restore highly-accurate DNA sequencing despite DNA damage. While it has been recognized that false mutations accumulate at fragment ends in duplex sequencing data due to the fill-in of short 5’ overhangs, the extent to which false mutations could manifest within the interior of each DNA duplex as a result of ER/AT has not been established. Our single-molecule sequencing assay has provided novel insight into ER/AT and mechanisms of DNA repair. Indeed, we were astonished to find that 7-9%, 15-17%, and 32-57% of base pairs >12 bp from the ends of each duplex in healthy cfDNA, cancer patient cfDNA, and FFPE tumor biopsies, respectively, could be resynthesized when conventional ER/AT methods were applied. Further, our induction of varied base and backbone damage has shown how the two together create the ‘perfect storm’ for errors when conventional ER/AT methods are applied. Our observation that both strand resynthesis and error rate increase with DNase I concentration suggests that the reliability of diagnostic tests such as liquid biopsies could be affected by the nuclease activity in an individual’s bloodstream. Given the wide variation in quality of clinical specimens, these findings have important implications for the field.

One limitation of our method to estimate fill-in via single-molecule sequencing is that it only uses two modified bases (d^6m^ATP and d^4m^CTP) which makes it challenging to pinpoint the exact base at which fill-in starts. Also, given the high error rates in single-molecule sequencing, we currently require multiple bases in a row with excess signal to detect fill-in and for the excess signal to be observed up to the 3’ end of the fragment. This means that we currently lack the resolution to resolve the fill-in of single nucleotides or short patches within the interior of a fragment. Additionally, while Duplex-Repair substantially limited resynthesis, there still appeared to be a small population of fragments with long fill-in, which could explain why errors in duplex sequencing remained. Yet, with our ability to measure strand resynthesis, we should be able to improve the method. Meanwhile, the observed fractions of bases resynthesized being highly correlated with sequencing error rates suggests that further limiting resynthesis may be able to maximize sequencing accuracy.

One limitation of Duplex-Repair is that resynthesis still occurs within gap regions and short (≤ 7 nt) 5’ overhangs after the DNA lesion repair and overhang removal step, as ExoVII cannot fully blunt 5’ overhangs. By first reducing the lengths of 5’ overhangs using ExoVII, it becomes possible to concentrate errors within fragment ends and filter against them *in silico* by their distance from fragment ends. However, our current strategy to induce strand breaks within gap regions bearing DNA damage is incomplete: first, we can neither account for all types of DNA lesions which may emerge, nor do we have enzymes available to correct all. There are also alterations involving canonical bases which, in the absence of a complementary strand, will be impossible to discern (e.g. deamination of 5-methylcytosine to produce thymine, or even insertions or deletions). Future strategies may involve digesting single-stranded DNA irrespective of whether it contains a recognizable lesion, or labelling resynthesized bases and excluding from analysis. Noteworthily, Abascal et al. recently reported nanorate sequencing (NanoSeq) that suppresses strand resynthesis during ER/AT by using a restriction enzyme to digest intact DNA to produce blunted dsDNA fragments and then non-A dideoxynucleotides during dA-tailing to block templated extension^31^. As a result, NanoSeq can achieve a reported error rate of < 5e-9 when applied to gDNA extracted from sperm and cord blood samples. This study further highlights the importance of limiting strand resynthesis for achieving high accuracy duplex sequencing. However, this method can only be applied to intact DNA. Furthermore, restriction enzyme digestion limits the coverage to ~ 30% of the human genome. An alternative method is to use mung bean nuclease to blunt fragmented DNA and then non-A dideoxynucleotides during dA-tailing. However, DNA fragments (containing nicks, gaps or overhangs) that are not fully blunted by mung bean nuclease will be rendered unusable for duplex sequencing.

Our study has shown that ER/AT methods function like a ‘pencil and eraser,’ rewriting the nucleobases downstream of discontinuities in the phosphodiester backbone, and spurring false detection of lesions or alterations originally confined to one strand. Meanwhile, our solution of Duplex-Repair offers one of the first known approaches to preserve the sequence integrity of duplex DNA and thus, improve the reliability of methods which leverage the duplicity of genetic information in DNA.

## DATA AVAILABILITY

All sequencing data generated in the course of this study will be deposited into a controlled access database such as dbGaP.

## ADDITIONAL INFORMATION

Supplementary information is available for this paper.

Correspondence and requests for materials should be addressed to V.A. Adalsteinsson.

## ACKNOWLEDGEMENTS

The authors would like to acknowledge the patients and their families for their contributions to this study. In addition, the authors acknowledge the Gerstner Family Foundation for its generous support.

## CONFLICT OF INTEREST

K. Xiong has a patent application filed with Broad Institute. T.R. Golub has advisor roles (paid) at Foundation Medicine, GlaxoSmithKline and Sherlock Biosciences. V.A. Adalsteinsson has a patent application filed with Broad Institute and is a member of the scientific advisory boards of AGCT GmbH and Bertis Inc., which were not involved in this study. The remaining authors report no conflicts of interest.

