## Supplementary Information for "Duplex-Repair enables highly accurate sequencing, despite DNA damage"

***Recalibrate capillary electrophoresis traces:***

Lengths of synthetic oligonucleotides are confirmed by IDT's mass spectrometry analysis (data not shown). However, the control peak locations reported from raw fragment analysis by using the Peak scanner 2 software differ from the expected positions (**Table S1**); the peak locations of 6-FAM tagged molecules consistently appear as underestimates whereas those with ATTO 550 present as overestimates.

To interpret the capillary electrophoresis data, we decide to recalibrate the peak locations by using a ladder of synthetic oligonucleotides with known lengths. **Equation S1-2** relates the oligonucleotide length to raw peak locations through linear regression.

$y=1.0381x-7.681$	Eq. S1

**Equation S1.** Linear regression of raw fragment analysis peak locations of the 6-FAM-tagged strands. Experimentally determined values for the oligos tagged with 6-FAM in the 100 bp, 90 bp, 80 bp and 70 bp ssDNA controls (**Table S1** oligos e, d, c, b respectively) were used to generate a model that relates actual oligonucleotide length (x) to the fragment analysis readout (y) for 6-FAM substrates (**Fig. S1A**).

$y=0.9666x+5.039$	Eq. S2

**Equation S2.** Linear regression of raw fragment analysis peak locations of the ATTO 550-tagged strands. Experimentally determined values for the oligos tagged with ATTO-550 in the 100 bp, 90 bp, 80 bp and 70 bp ssDNA controls (**Table S1** oligos i, h, g, f respectively) were used to generate a model that relates actual oligonucleotide length (x) to the fragment analysis readout (y) for ATTO-550 substrates (**Fig. S1B**).

***Quantification of library conversion efficiency by ddPCR:***

To quantify library conversion efficiency, a ddPCR assay was designed to target the flanking adapter regions. Only fragments with successful double ligation were exponentially amplified within the QX200 ddPCR EvaGreen Supermix (Bio-Rad) and thus detected.

*ddPCR assay design*

Primer 1: CACTCTTTCCCTACACGACG

Primer 2: AGTTCAGACGTGTGCTCTTC

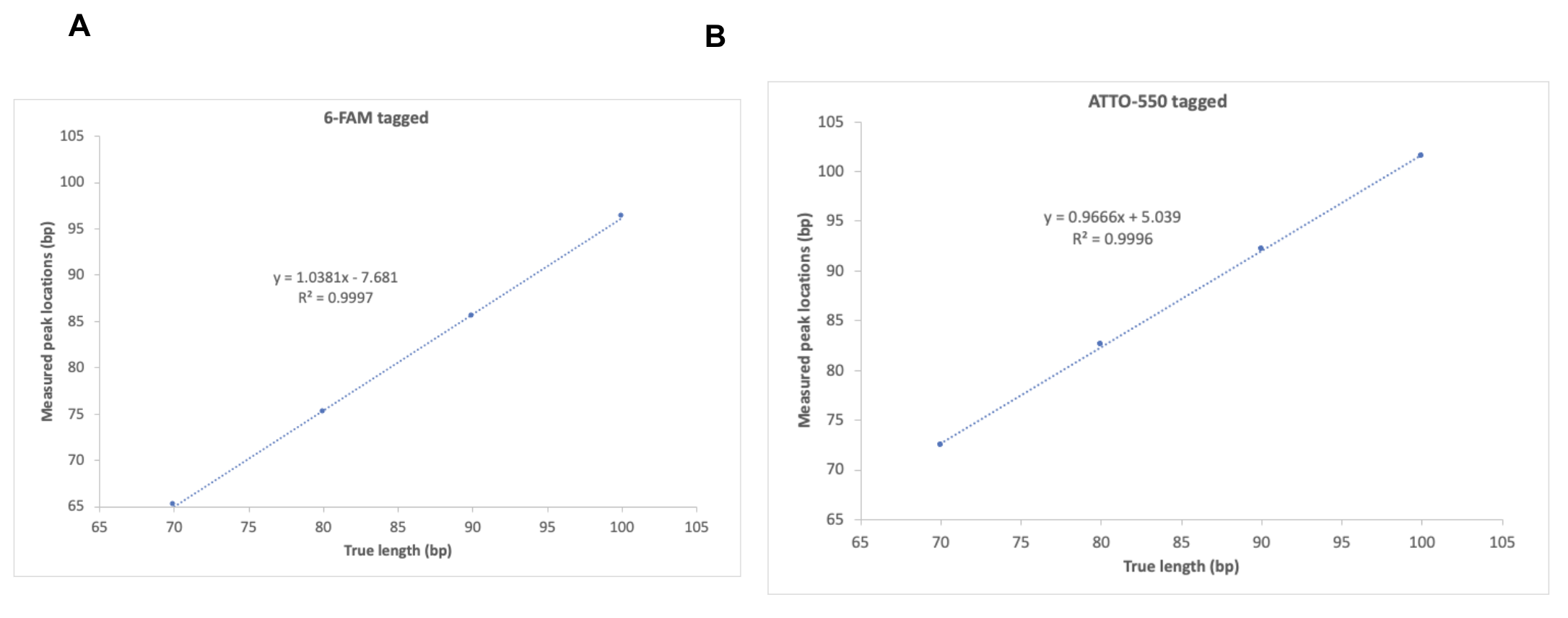

**Figure S1: Linear regression of measured capillary electrophoresis peak locations vs. true lengths for (a) 6-FAM-tagged and (b) ATTO-550 tagged oligonucleotides**. True lengths of oligonucleotides are confirmed by IDT’s mass spectrometry analysis (data not shown).

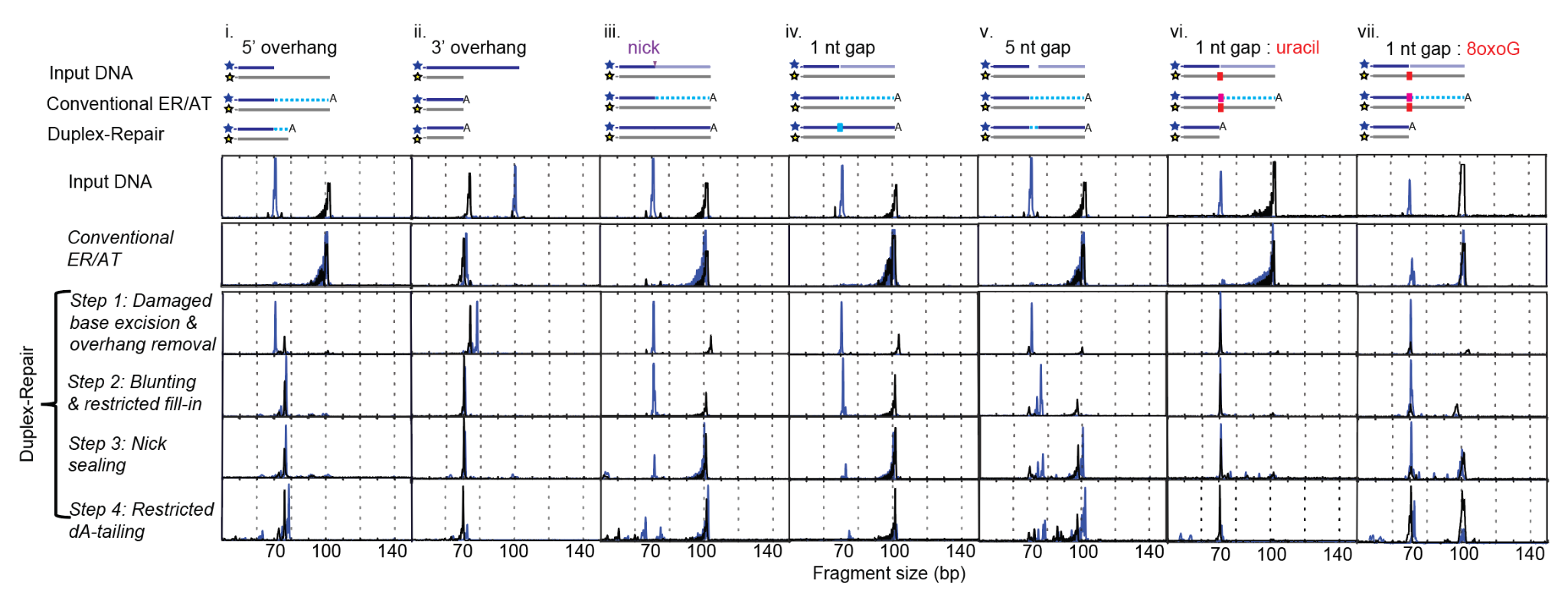

**Figure S2: Capillary electrophoresis analysis of synthetic duplexes subjected to each step of Duplex-Repair, versus conventional ER/AT.** Each step of duplex repair imparts its intended functionality in producing the intended major product as depicted in Fig. 1 to minimize strand resynthesis seen with Conventional ER/AT. Oligonucleotides with a (i) 5’ overhang, (ii) 3’overhang, (iii) nick, (iv) 1 nucleotide gap, (v) 5 nucleotide gap, (vi) uracil across from a 1 nucleotide gap, and (vii) 8oxoG across from a 1 nucleotide gap were subjected to conventional ER/AT and each step of Duplex Repair and sent for capillary electrophoresis. The top strand of each oligonucleotide is labelled with 6-FAM on the 5’ end, and the fragment size distributions following each treatment are represented by blue curves. The bottom strand of each oligonucleotide is labelled with ATTO-550 on the 3’ end, and the fragment size distributions following each treatment are represented by black curves.

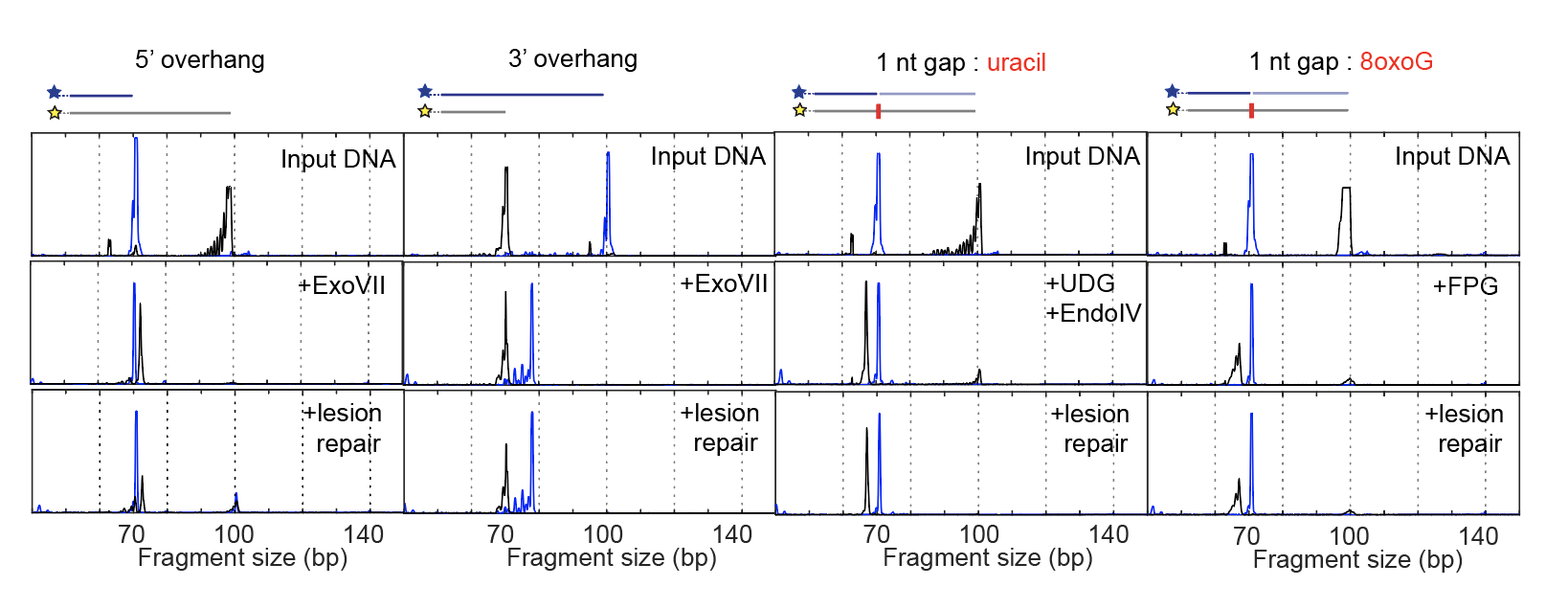

**Figure S3: Characterization of the activity of key enzymes in the lesion repair enzyme cocktail by capillary electrophoresis.** The activity of key enzymes to rectify each damage motif (middle) is not impacted by other enzymes in the lesion repair enzyme cocktail (bottom). The “lesion repair” condition indicates treatment with Endonuclease IV (EndoIV), Formamidopyrimidine [fapy]-DNA glycosylase (Fpg), Uracil-DNA glycosylase (UDG), T4 pyrimidine DNA glycosylase (T4 PDG), and Endonuclease VIII (EndoVIII), and Exonuclease VII (ExoVII).

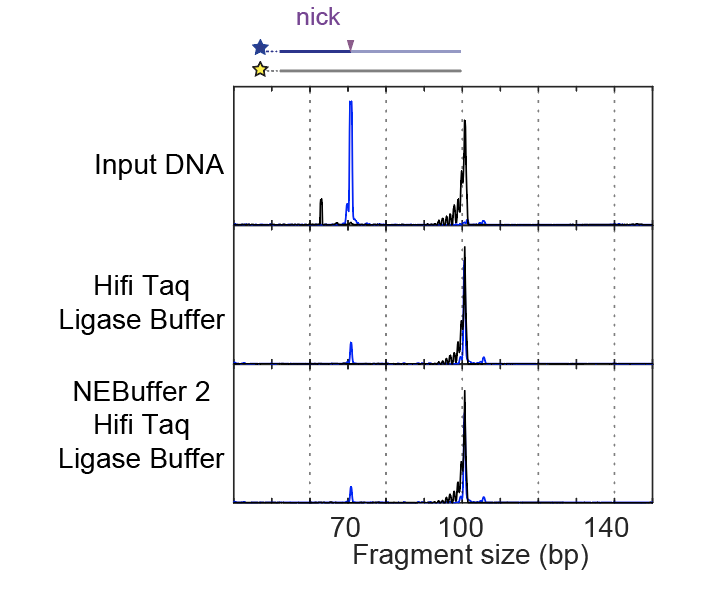

**Figure S4: Characterization of the activity of HiFi Taq DNA ligase by capillary electrophoresis.** HiFi Taq DNA ligase efficiently seals nicks in NEBuffer 2 and HiFi Taq ligase buffer mix (bottom) as it does in HiFi Taq ligase buffer alone (middle).

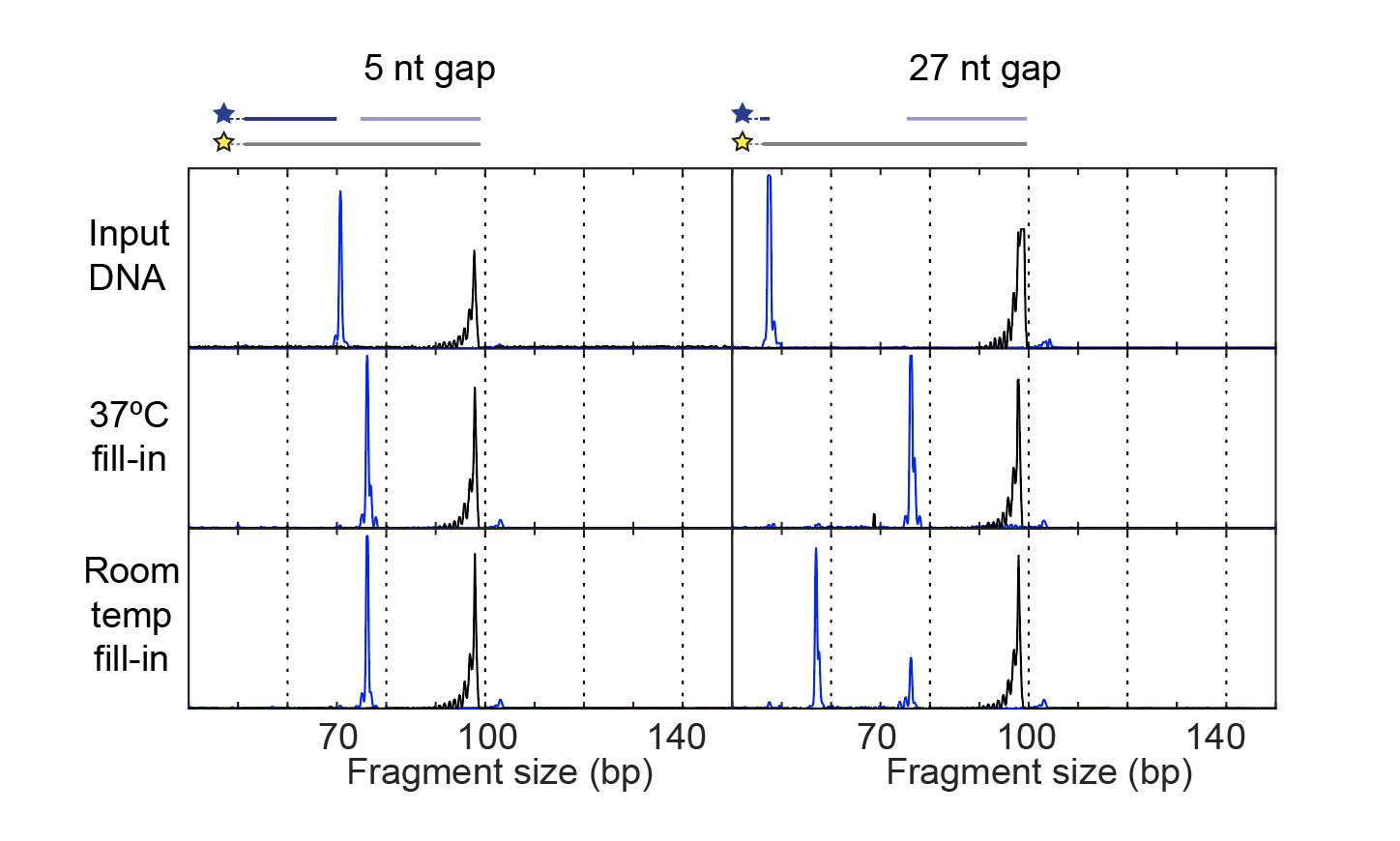

**Figure S5: Characterization of the activity of T4 DNA polymerase and T4 polynucleotide kinase by capillary electrophoresis.** T4 DNA polymerase efficiently fills in 5 or 27 nt gaps at 37 ^o^C in NEBuffer 2 with no detectable strand-displacement activity (middle). The efficiency of T4 DNA polymerase filling in 27 nt gaps at room temperature, however, is significantly lower (bottom).

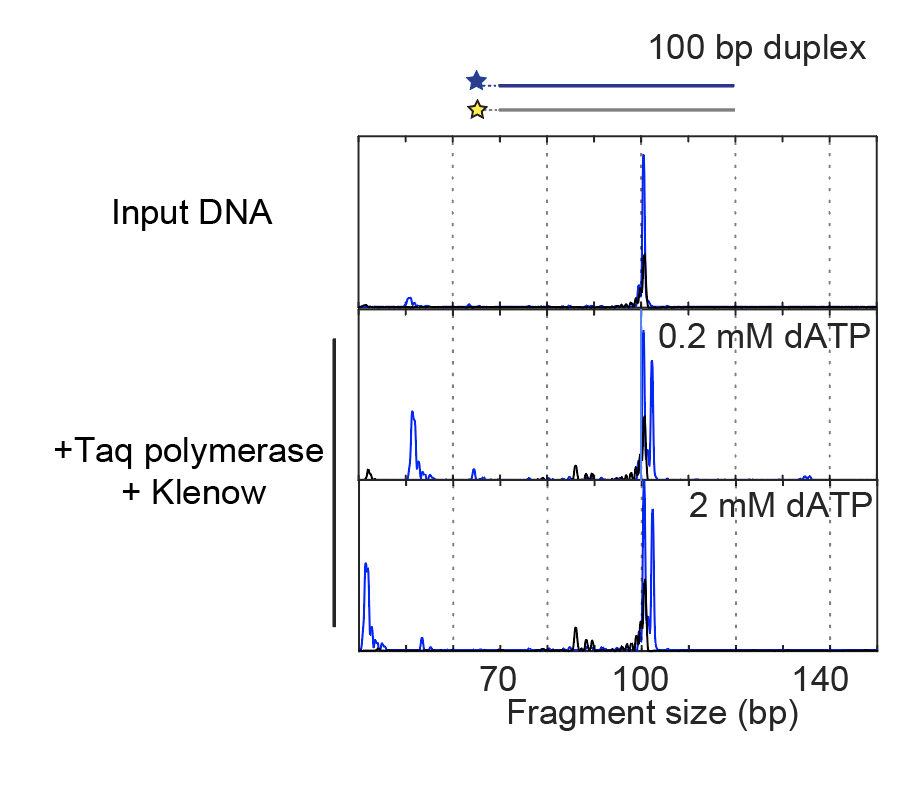

**Figure S6: Characterization of the activity of Klenow fragment (exo-) and Taq DNA polymerase by capillary electrophoresis.** Klenow (exo-) and Taq DNA polymerase efficiently perform dA-tailing with only dATP present at concentrations of 0.2 mM (middle) or 2 mM (bottom).

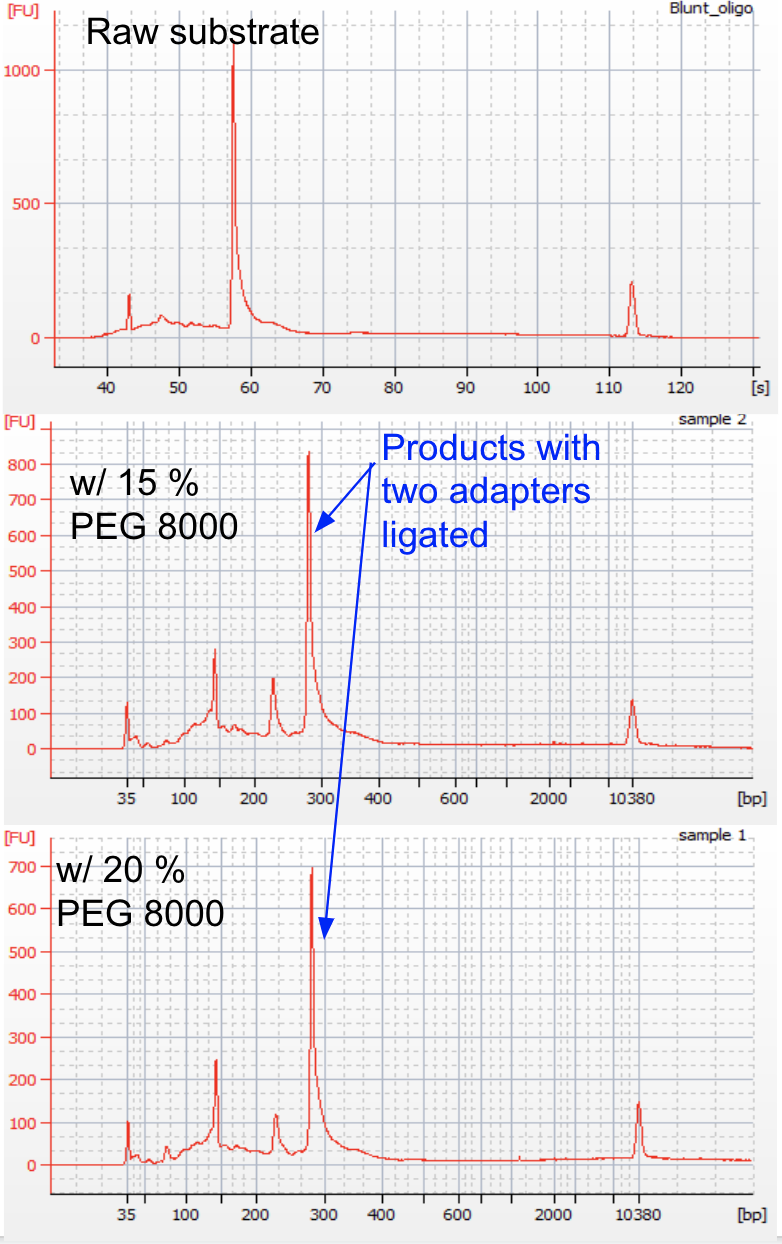

**Figure S7: Characterization of the activity of T4 DNA ligase and 5' deadenylase by BioAnalyzer.** T4 DNA ligase and 5' deadenylase efficiently ligate NGS adapters to a 166 bp blunted duplex with dA tails in the presence of 15 (top) or 20% (bottom) weight by volume (w/v) PEG 8000. To minimize spurious intermolecular ligation at high PEG concentrations, Duplex-Repair only uses 10% w/v PEG 8000 during adapter ligation. Of note: the unit of the x axis of the top panel could not be converted to bp by BioAnalyzer software.

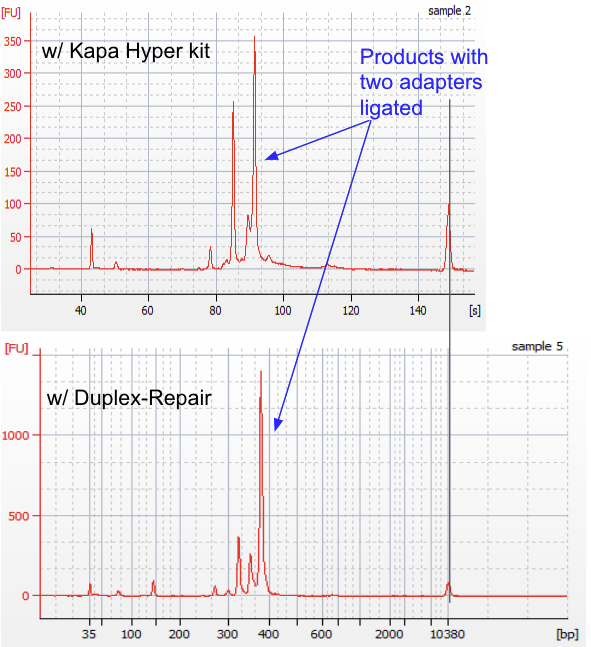

**Figure S8: Characterization of the combined efficiency of dA-tailing and adapter ligation by BioAnalyzer.** The combined efficiency of dA-tailing and adapter ligation of Duplex-Repair could be higher than that of the Kapa Hyper kit. The input was a 274 bp blunted duplex. Of note, the unit of the x axis of the top panel could not be converted to bp by BioAnalyzer software.

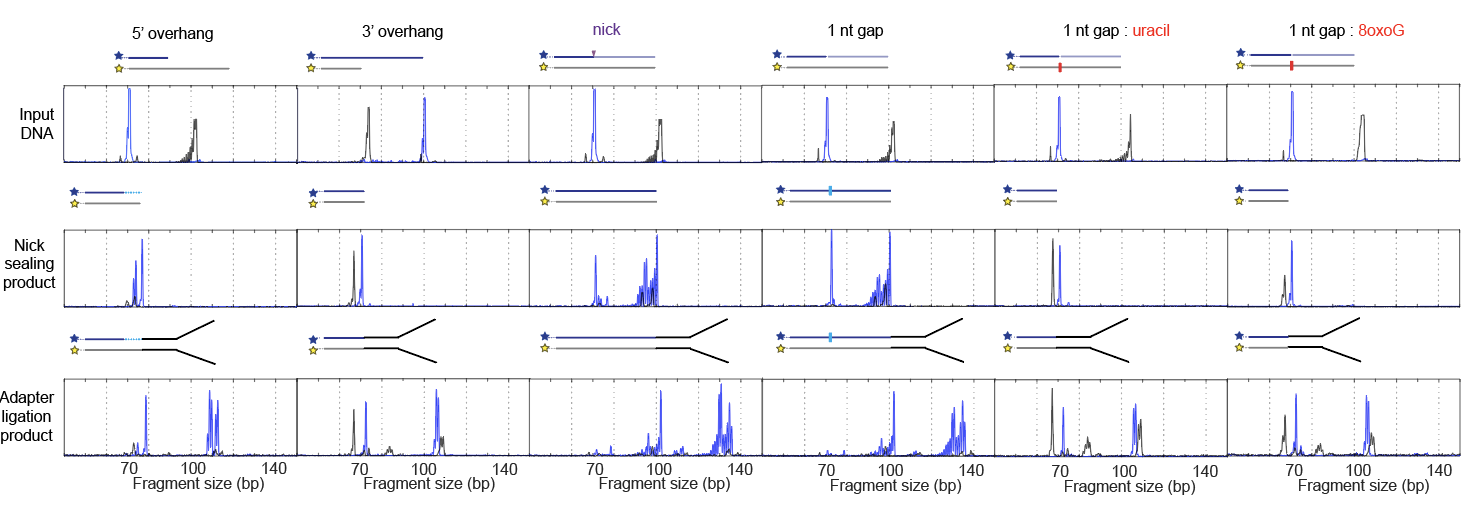

**Figure S9: Characterization of the performance of Duplex-Repair (after optimizing reaction conditions and eliminating multiple Ampure cleanups) by capillary electrophoresis**. Duplex-Repair facilitates the formation of a major product of NGS adapter-ligated oligonucleotides that are ready for sequencing applications. The ‘nick sealing products’ (middle) were collected following steps 1-3 of duplex repair but prior to dA-tailing. The ‘adapter ligated products’ (bottom) have undergone the entire Duplex-Repair protocol and ligation to NGS adapters, which add an additional 39-40 or 37-38 bp (unique molecular indices can be either 3 or 4 base pairs) to the exposed 3’ and 5’ ends of oligonucleotides after Duplex-Repair respectively (note: adapters in schematic not drone to scale).

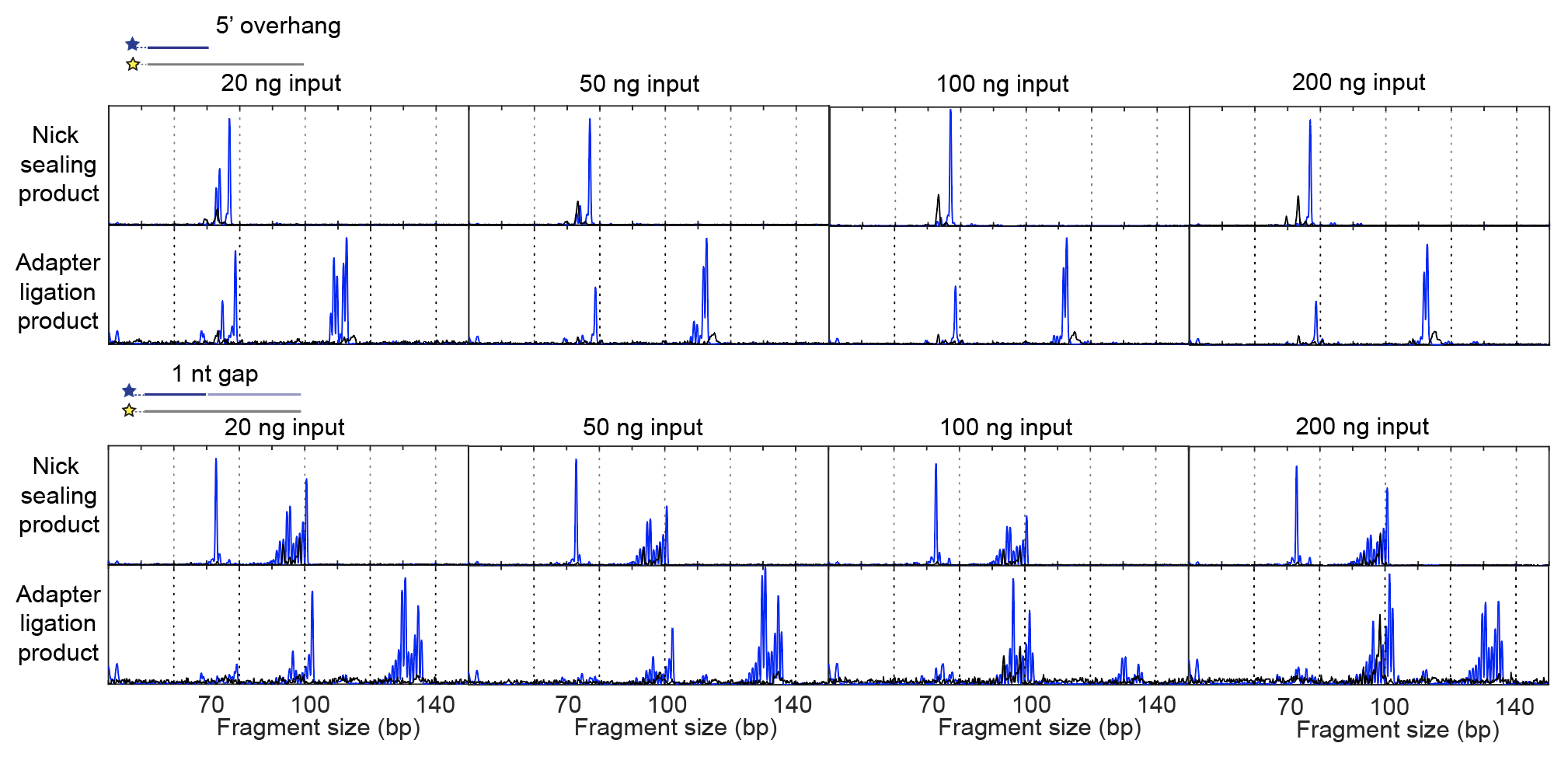

**Figure S10: Characterization of the performance of Duplex-Repair (after optimizing reaction conditions and eliminating multiple Ampure cleanups) as a function of DNA input mass by capillary electrophoresis.** Duplex-Repair is effective at preparing cfDNA inputs ranging from 20 to 200 ng for NGS. The ‘nick sealing products’ (top rows) were collected following steps 1-3 of duplex repair but prior to dA-tailing. The ‘adapter ligated products’ (bottom rows) have undergone the entire Duplex-Repair protocol and ligation to NGS adapters, which add an additional 39-40 or 37-38 bp (unique molecular indices can be either 3 or 4 base pairs) to the exposed 3’ and 5’ ends of oligonucleotides after Duplex-Repair respectively.

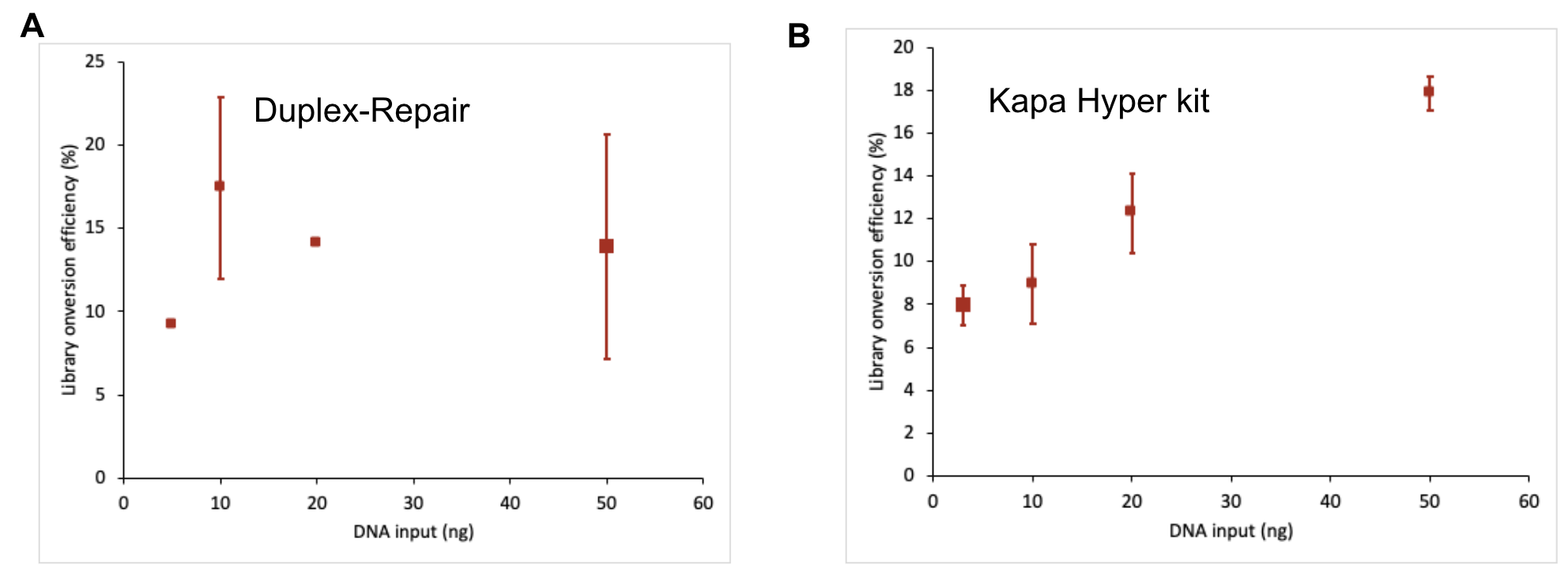

**Figure S11: The measured library conversion efficiencies of Duplex-Repair vs. the Kapa Hyper kit as a function of a gDNA input by using a ddPCR assay.** The library conversion efficiencies of Duplex-Repair are comparable to this with conventional ER/AT using the Kapa Hyper kit. ddPCR primers used are detailed in the Supplementary Text.

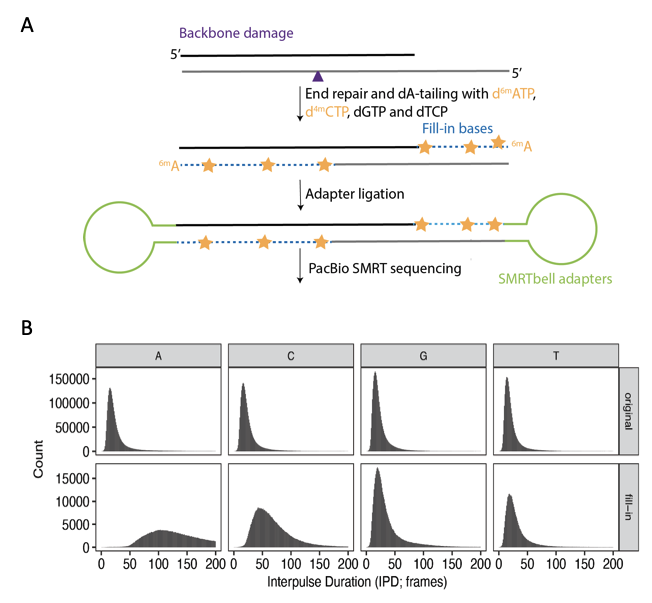

**Figure S12: Establishment of an assay for quantifying the number of bases resynthesized during ER/AT.** Histogram of aggregate bases and their IPDs, labeled as original or fill-in based on which region of the synthetic oligos they were derived from. Regions that divide original and fill-in regions were avoided for collection.

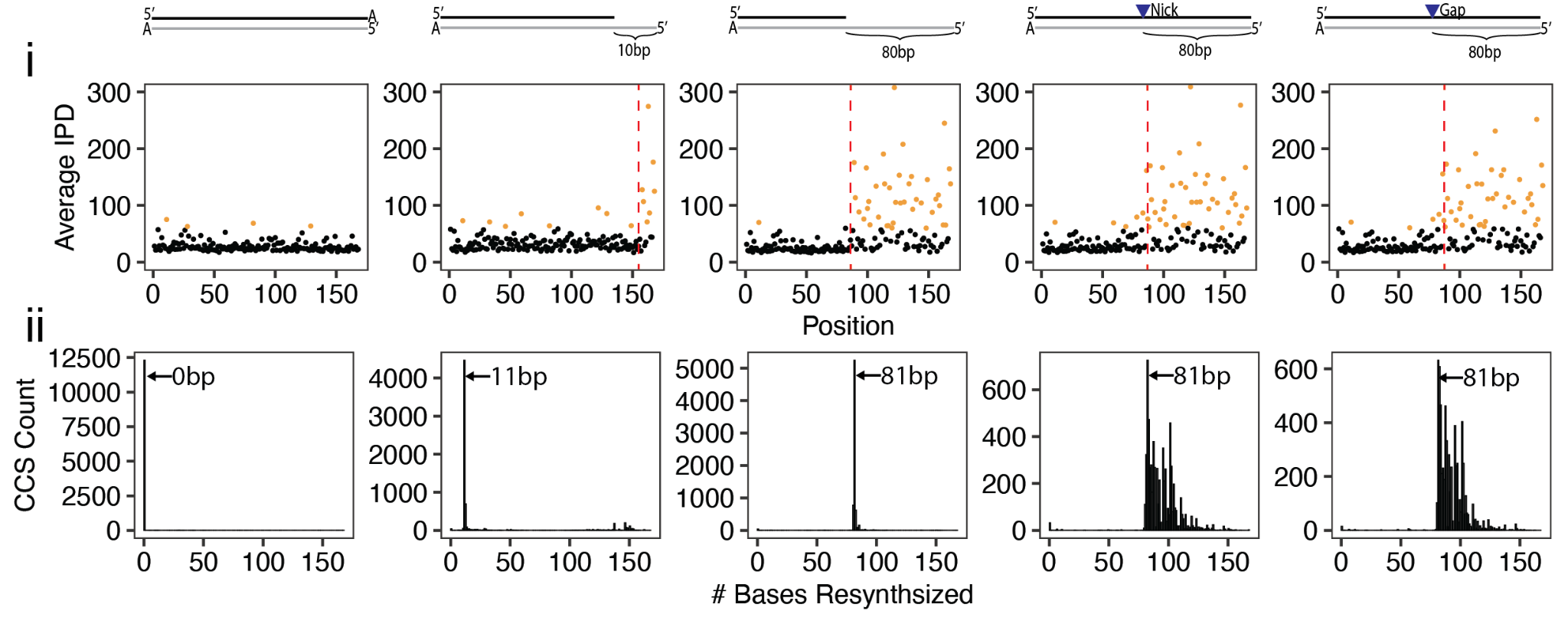

**Figure S13:** Measured interpulse duration (IPD; in frames) (i) and predicted percentage of bases resynthesized (ii) as a function of the base position on five synthetic oligonucleotides treated with conventional ER/AT and with modified dNTPs. Longer IPDs, colored orange if greater than 60 frames, result from modified bases. Red dashed lines indicate where resynthesis is expected to start during ER/AT.

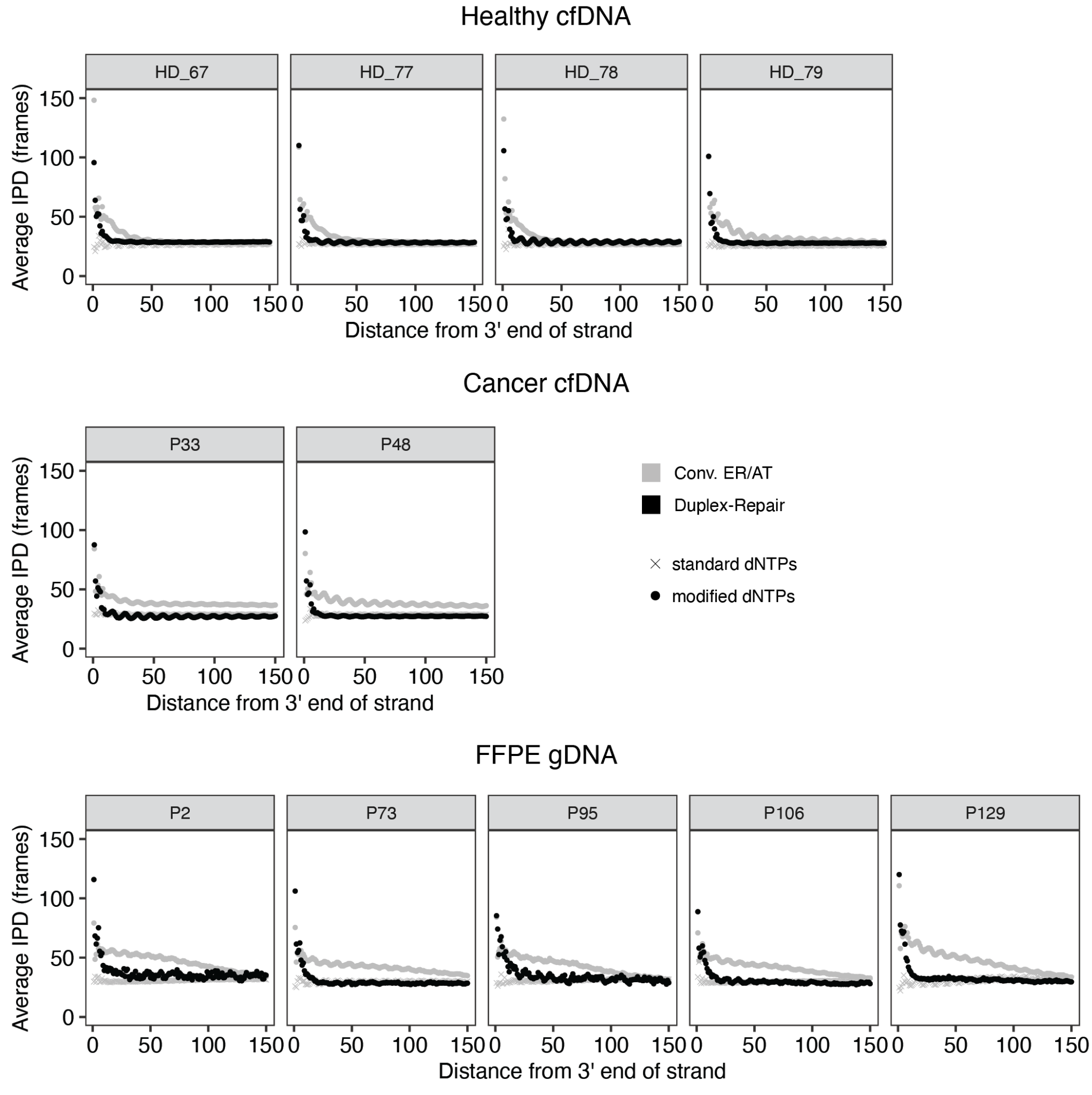

**Figure S14: Identification of resynthesized regions by reading out interpulse durations during PacBio SMRT sequencing.** Average interpulse durations (IPD; in frames) for each position relative to distance from the end of the original duplex DNA fragment for healthy cfDNA, cancer patient cfDNA, and FFPE tumor biopsies across several individuals.

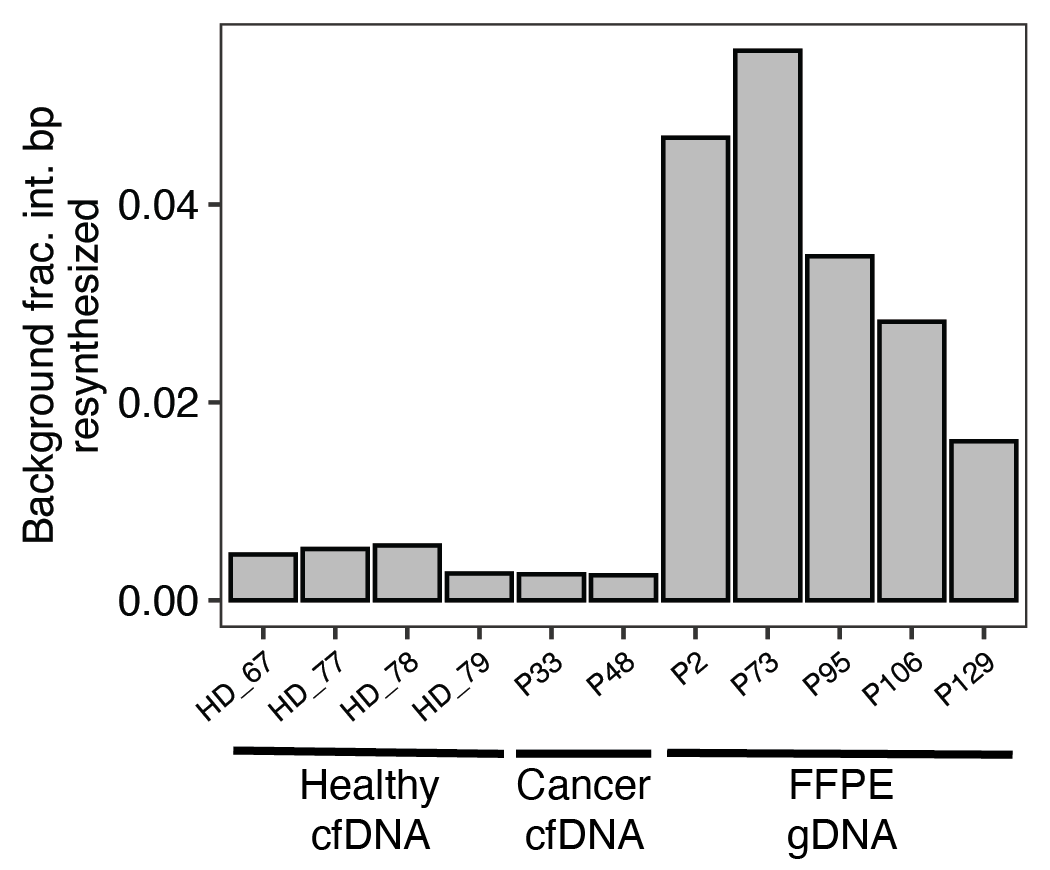

**Figure S15: Background estimated resynthesis of interior base pairs using standard dNTPs across FFPE and cfDNA sample types.**

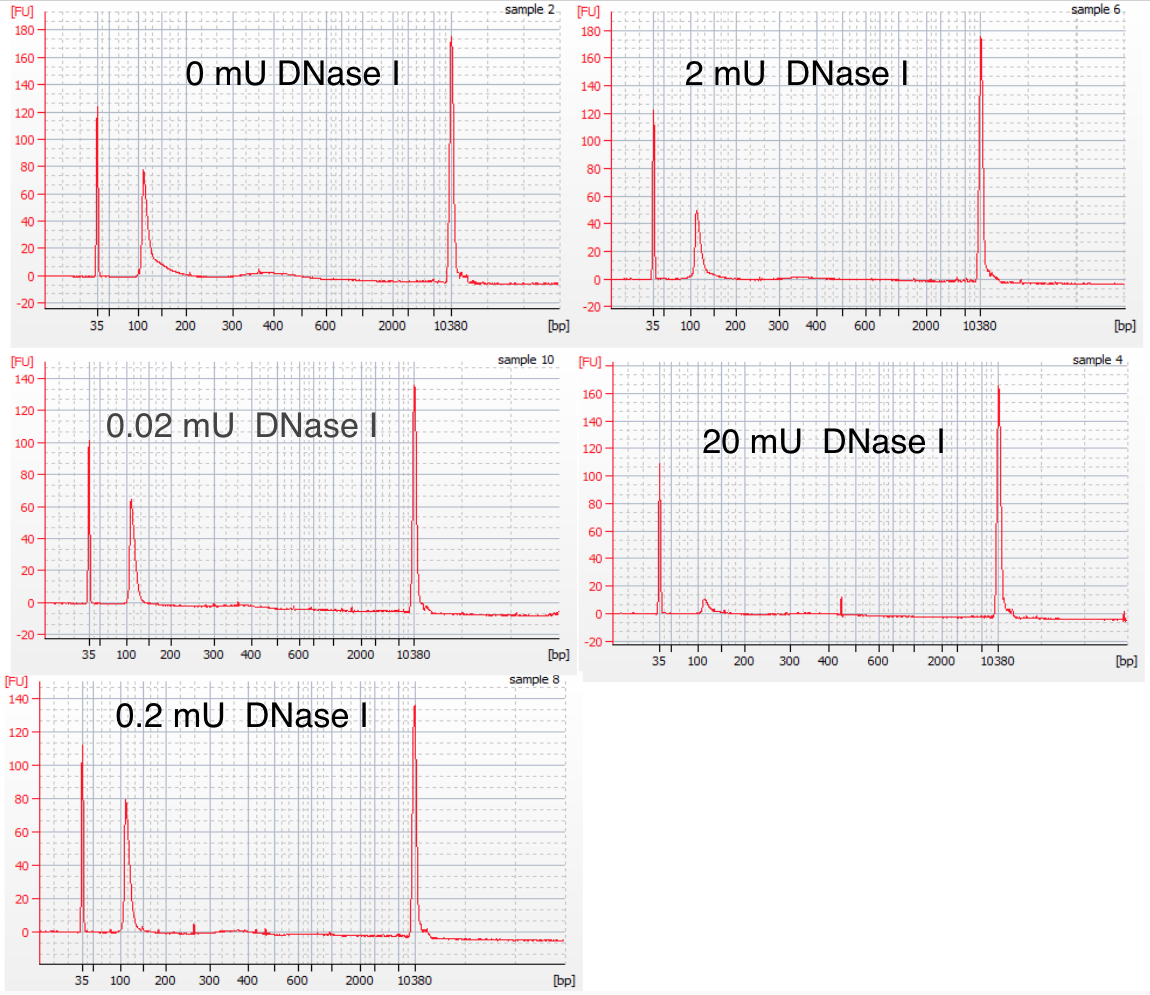

**Figure S16: Characterization of the activity of DNase 1 by BioAnalyzer**. The input was a 100 bp dsDNA oligo. The results show that up until 20 mU of DNase 1, the dominant fragment length is still 100 bp.

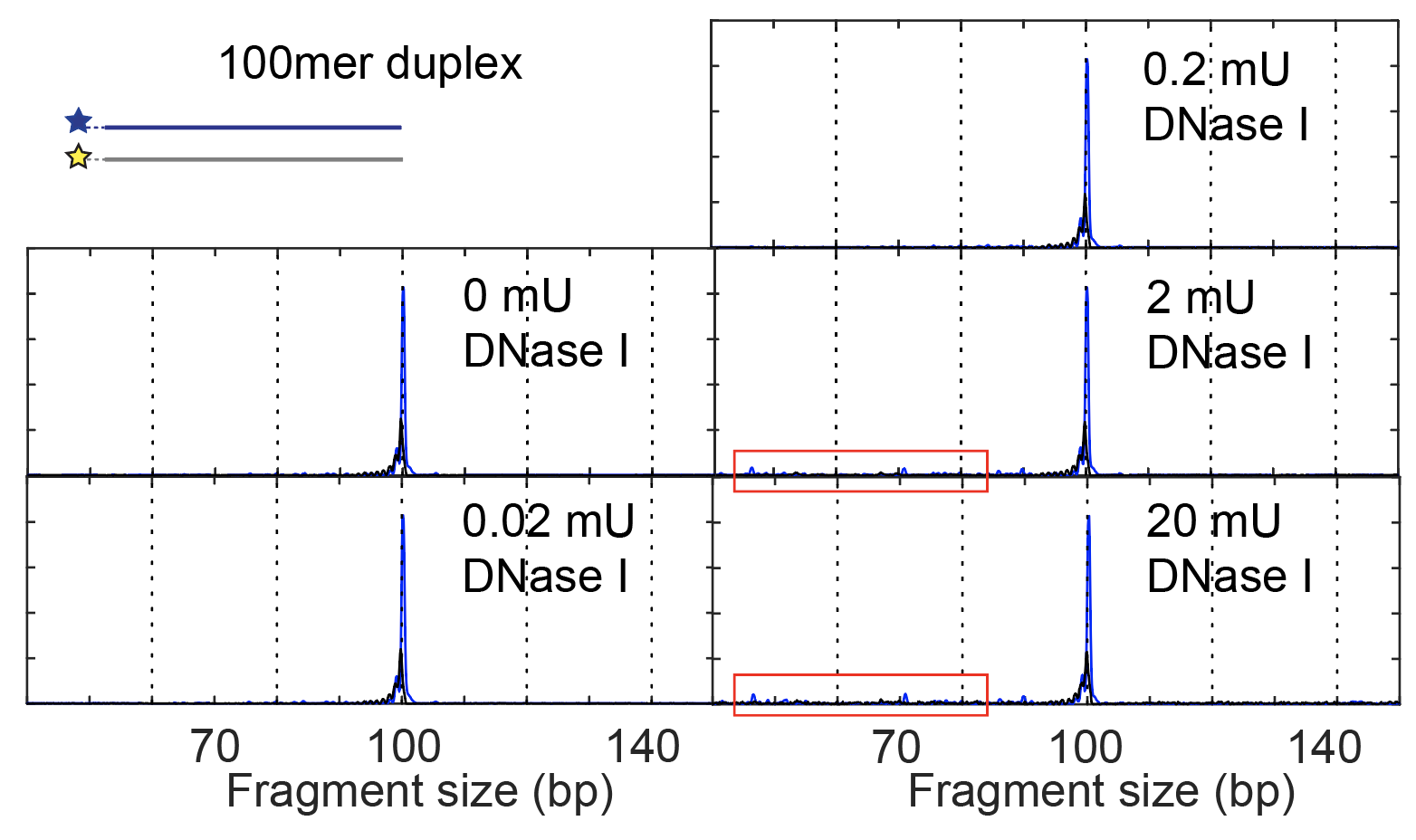

**Figure S17: Characterization of the activity of DNase 1 by capillary electrophoresis**. For all concentrations of DNAse 1 tested, the major product as determined by capillary electrophoresis is the 100mer duplex. However, intermediate-sized fragments (highlighted in red boxes) are detected with 2 and 20 mU of DNase 1 , suggesting that ≥2 mU of DNase 1 nick but do not significantly degrade dsDNA. These intermediate-sized fragments are present in capillary electrophoresis traces, as heat pretreatment and denaturation is required, but not on BioAnalyzer traces in which there is no denaturation (Fig. S15).

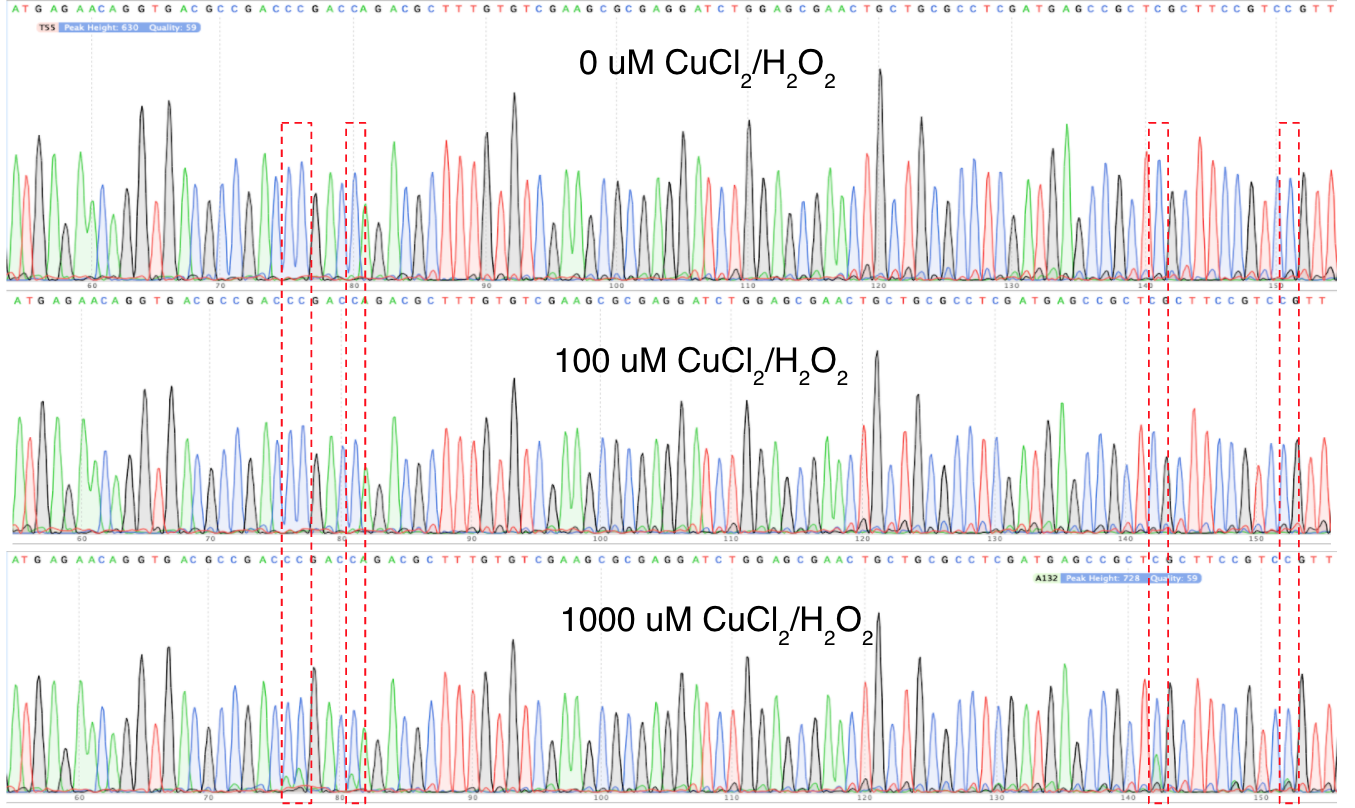

**Figure S18: Characterization of the oxidation activity of CuCl_2_/H_2_O_2_ by Sanger sequencing.** The input was a 274 bp dsDNA oligo and was treated with different concentrations of CuCl_2_/H_2_O_2_. The red boxes indicate where C->A mutations are detected when treated with 1000 uM CuCl_2_/H_2_O_2_.

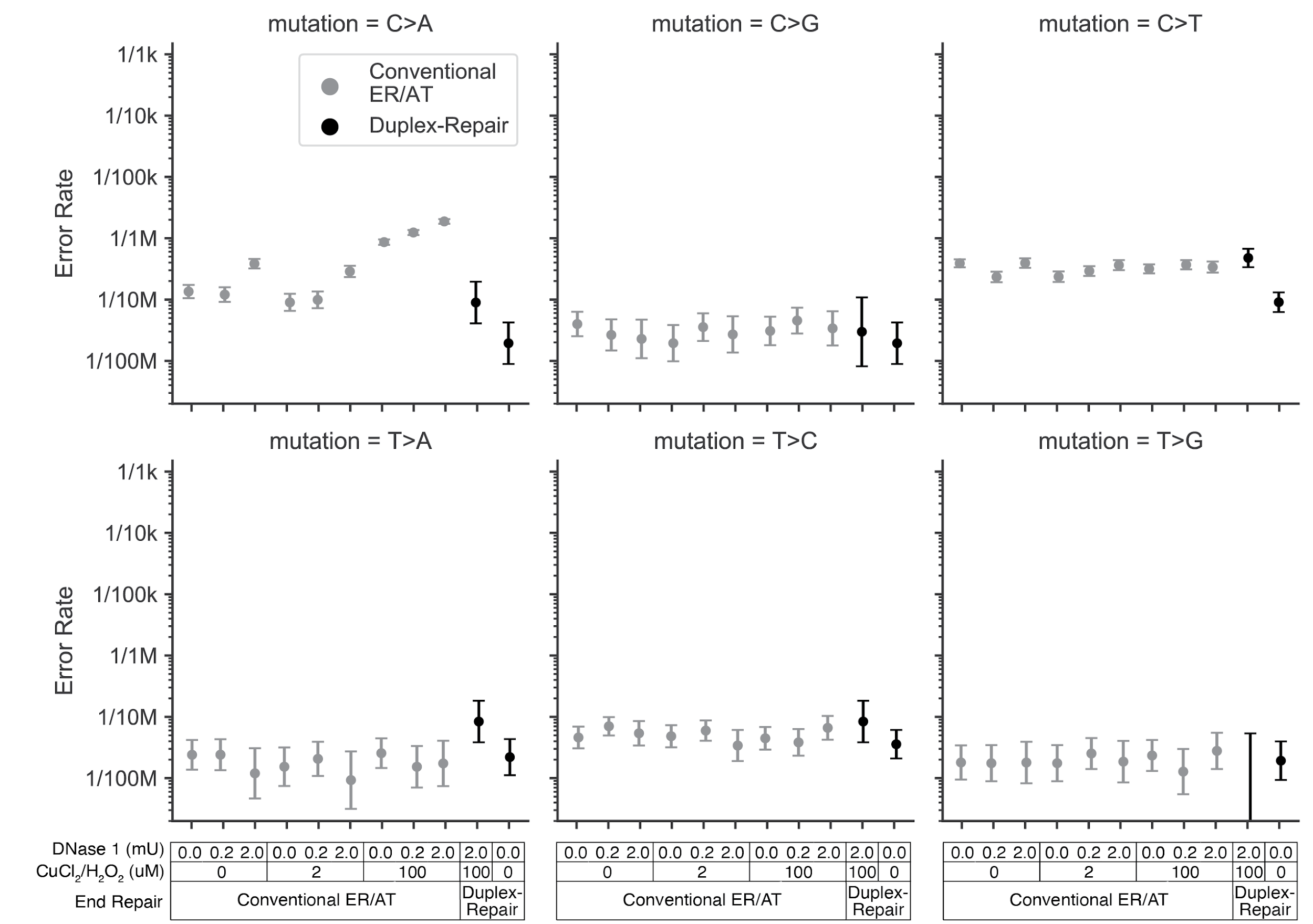

**Figure S19: Error rates by mutation context observed in healthy donor cfDNA treated with varied concentrations of CuCl_2_/H_2_O_2_ and DNase I.**

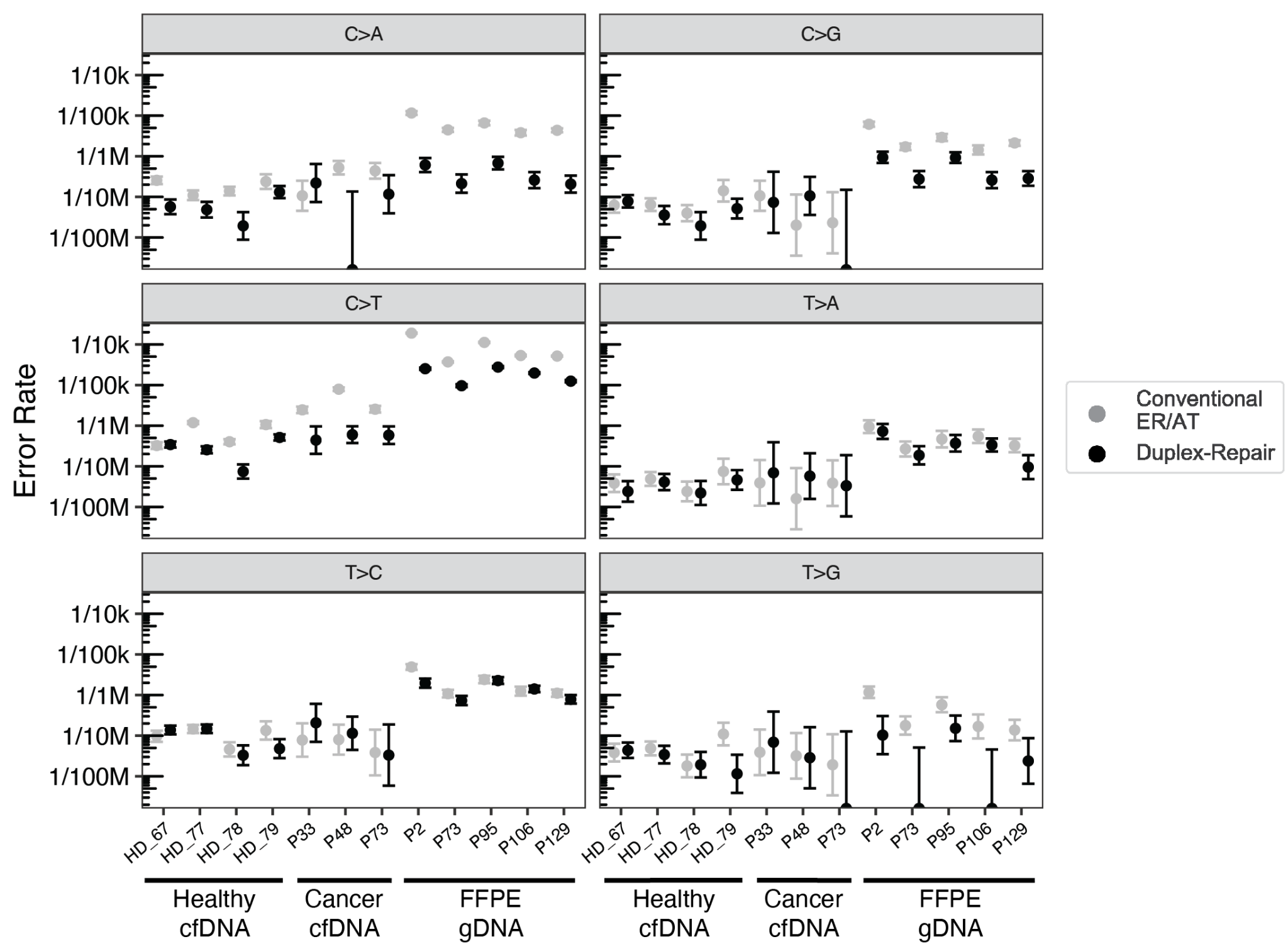

**Figure S20: Error rates by mutation context observed in duplex sequencing of a pan-cancer panel for cfDNA samples and FFPE tumor biopsies treated with conventional ER/AT vs. Duplex-Repair.**

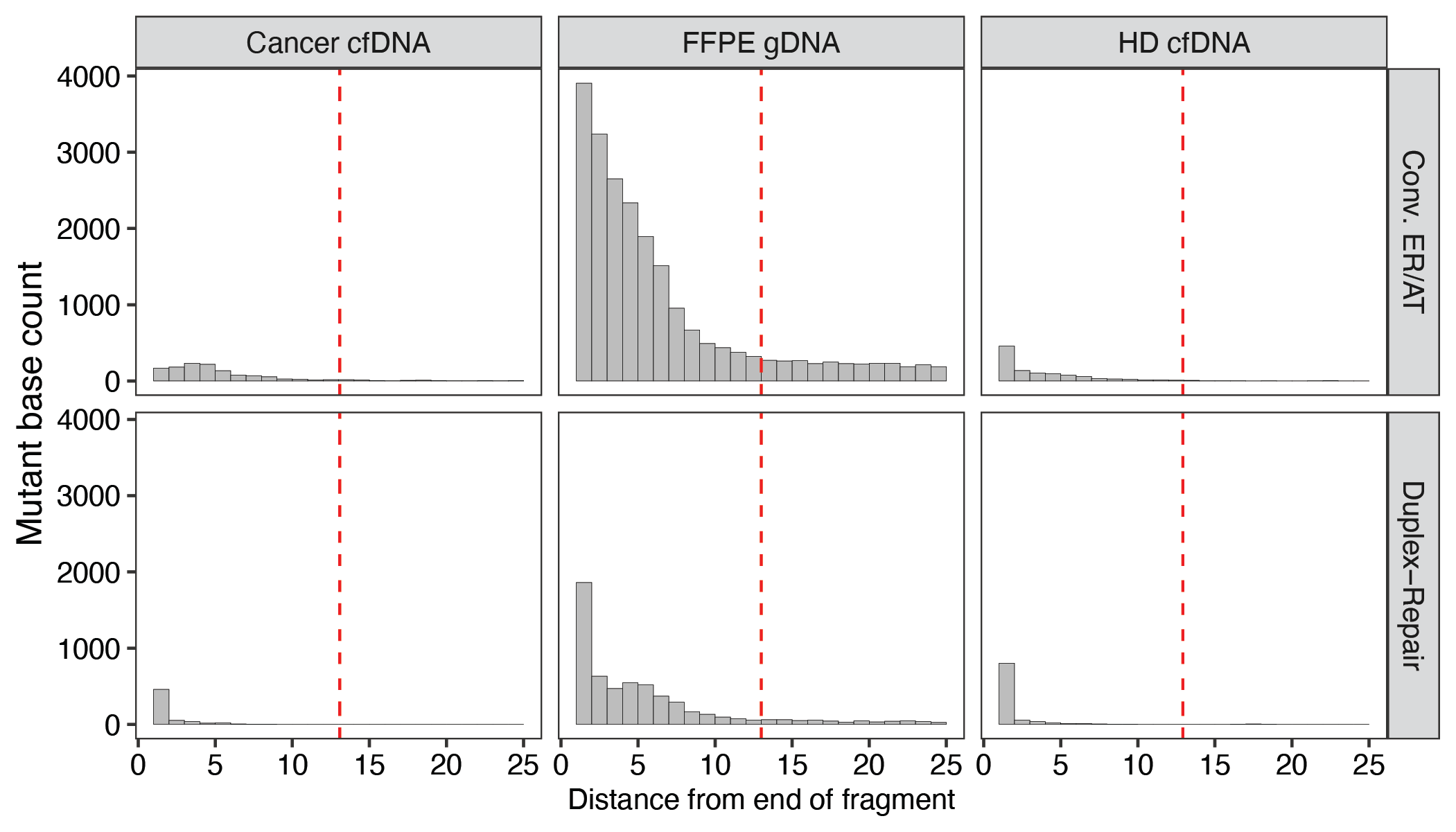

**Figure S21: Distance of mutant duplex bases from closest DNA fragment end for cfDNA collected from healthy donors and cancer patients as well as gDNA from FFPE tumor biopsies. Samples underwent either conventional ER/AT or Duplex-Repair.**

**Table S1: DNA sequences of synthetic oligonucleotides used in this study**. Asterisks (*) indicate the presence of a C3 spacer or phosphorothioate bonds that protect fluorophores from being cleaved by nucleases.

| Oligo ID | Fluorophore end | Fluorophore | Sequence | Length (bp) |
| --- | --- | --- | --- | --- |
| a | 5' | 6-FAM | 6FAM//iSpC3/GCGTCACCAGCCACGCGAGCCGGATGAGGATCCGTGACGCGAAGT | 48 |
| b | 5' | 6-FAM | 6FAM//iSpC3/GCGTCACCAGCCACGCGAGCCGGATGAGGATCCGTGACGCGAAGTCCTGGTACCGCCGCTCGCTTCCGAC | 70 |
| c | 5' | 6-FAM | 6FAM//iSpC3/GCGTCACCAGCCACGCGAGCCGGATGAGGATCCGTGACGCGAAGTCCTGGTACCGCCGCTCGCTTCCGACCGGTTCTCCA | 80 |
| d | 5' | 6-FAM | 6FAM//iSpC3/GCGTCACCAGCCACGCGAGCCGGATGAGGATCCGTGACGCGAAGTCCTGGTACCGCCGCTCGCTTCCGACCGGTTCTCCACCGAGCGACC | 90 |
| e | 5' | 6-FAM | 6FAM//iSpC3/GCGTCACCAGCCACGCGAGCCGGATGAGGATCCGTGACGCGAAGTCCTGGTACCGCCGCTCGCTTCCGACCGGTTCTCCACCGAGCGACCTAATATTAAT | 100 |
| f | 3' | ATTO 550 | GTCGGAAGCGAGCGGCGGTACCAGGACTTCGCGTCACGGATCCTCATCCGGCTCGCGTGGCTGGTGACGC/iSpC3//3ATTO550 | 70 |
| g | 3' | ATTO 550 | TGGAGAACCGGTCGGAAGCGAGCGGCGGTACCAGGACTTCGCGTCACGGATCCTCATCCGGCTCGCGTGGCTGGTGACGC/iSpC3//3ATTO550 | 80 |
| h | 3' | ATTO 550 | GGTCGCTCGGTGGAGAACCGGTCGGAAGCGAGCGGCGGTACCAGGACTTCGCGTCACGGATCCTCATCCGGCTCGCGTGGCTGGTGACGC/iSpC3//3ATTO550 | 90 |
| i | 3' | ATTO 550 | ATTAATATTAGGTCGCTCGGTGGAGAACCGGTCGGAAGCGAGCGGCGGTACCAGGACTTCGCGTCACGGATCCTCATCCGGCTCGCGTGGCTGGTG*A*C*G*C/3ATTO550 (*phosphorothioate bonds) | 100 |
| j | 3' | ATTO 550 | ATTAATATTAGGTCGCTCGGTGGAGAACC**/i8oxodG**/GTCGGAAGCGAGCGGCGGTACCAGGACTTCGCGTCACGGATCCTCATCCGGCTCGCGTGGCTGGTGA*C*G*C*/3ATTO550N/ (*phosphorothioate bonds) | 100 |
| k | 3' | ATTO 550 | ATTAATATTAGGTCGCTCGGTGGAGAACC**U**GTCGGAAGCGAGCGGCGGTACCAGGACTTCGCGTCACGGATCCTCATCCGGCTCGCGTGGCTGGTGA*C*G*C*/3ATTO550N/ (*phosphorothioate bonds) | 100 |
| l | / | / | CGGTTCTCCACCGAGCGACCTAATATTAAT | 30 |
| m | / | / | GGTTCTCCACCGAGCGACCTAATATTAAT | 29 |
| n | / | / | CTCCACCGAGCGACCTAATATTAAT | 25 |
| o | / | / | GTCAAGGGTAATGGACAGTAGGTGTGGTGGAACATACTTCCAGCACTCAAGAAGCTGAAGCAGGCAGATCTCTGTCAGTTCATGACCACTGCTGTCTACATGGTGAGCTCCAAGCCAGCCAGGCAAGAAGTGACACTCAGGtCTCGCATTGCTcagACGgCaggcA | 166 |
| p | / | / | TgcctGcCGTctgAGCAATGCGAGaCCTGAGTGTCACTTCTTGCCTGGCTGGCTTGGAGCTCACCATGTAGACAGCAGTGGTCATGAACTGACAGAGATCTGCCTGCTTCAGCTTCTTGAGTGCTGGAAGTATGTTCCACCACACCTACTGTCCATTACCCTTGACA | 166 |
| q | / | / | GTCAAGGGTAATGGACAGTAGGTGTGGTGGAACATACTTCCAGCACTCAAGAAGCTGAAGCAGGCAGATCTCTGTCAGTTCATGACCACTGCTGTCTACATGGTGAGCTCCAAGCCAGCCAGGCAAGAAGTGACACTCAGGtCTCGCATTGCTcag | 156 |
| r | / | / | GTCAAGGGTAATGGACAGTAGGTGTGGTGGAACATACTTCCAGCACTCAAGAAGCTGAAGCAGGCAGATCTCTGTCAGTTCATGAC | 86 |
| s | / | / | CACTGCTGTCTACATGGTGAGCTCCAAGCCAGCCAGGCAAGAAGTGACACTCAGGtCTCGCATTGCTcagACGgCaggcA | 80 |
| t | / | / | ACTGCTGTCTACATGGTGAGCTCCAAGCCAGCCAGGCAAGAAGTGACACTCAGGtCTCGCATTGCTcagACGgCaggcA | 79 |

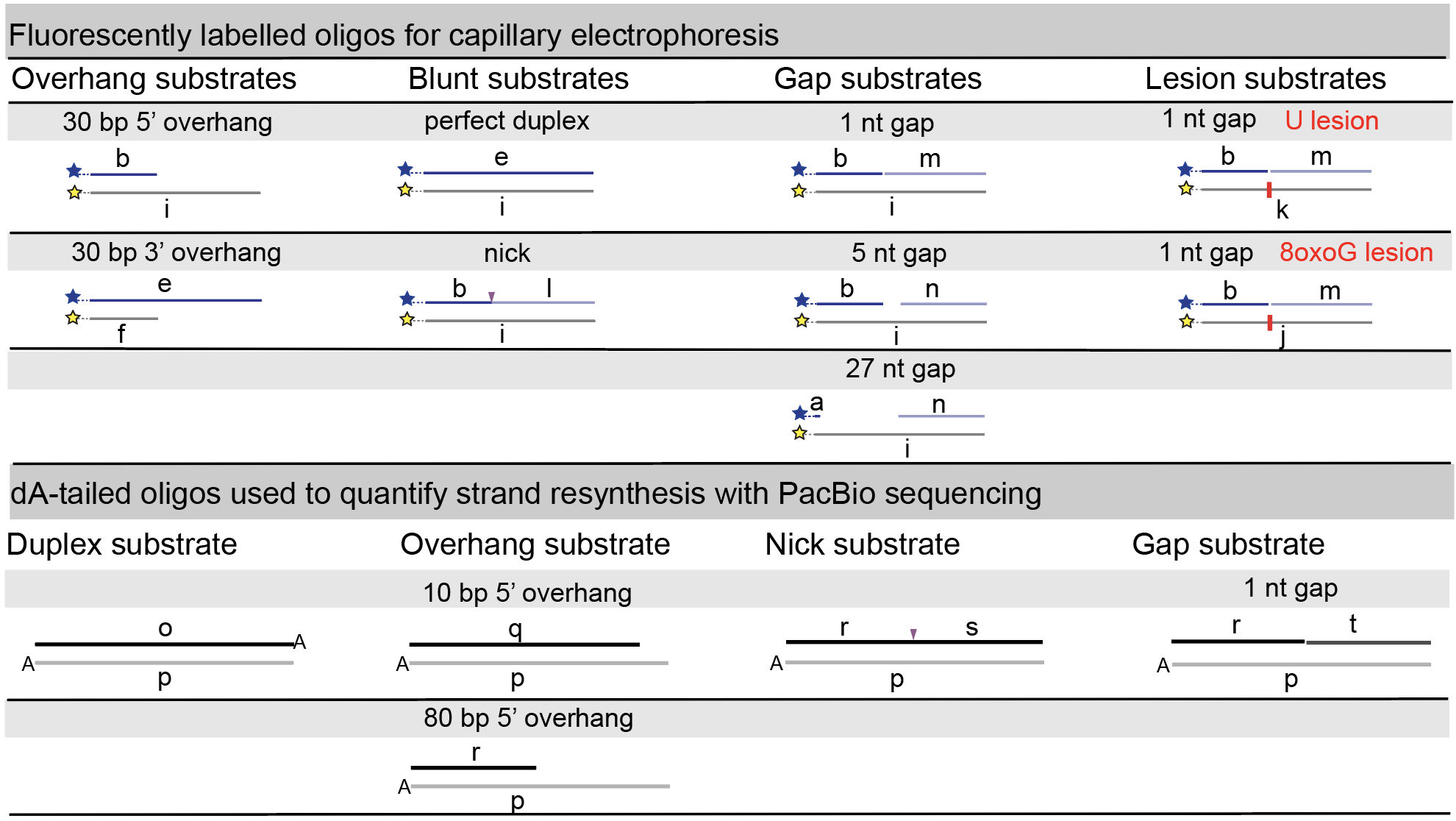

**Table S2: Quantification of DNA loss after DNase 1 treatment**. The input was 20 ng of a 100 bp dsDNA oligo. *the low yield indicates a significant loss during the Ampure bead cleanup step; ** the concentration of the 2nd biological replicate is below the detection limit of the Qubit assay.

| Dnase I amount (mU) | Yields of DNA products after DNase 1 treatment, mean ± SD (with two biological replicates) |
| --- | --- |
| 0 | 2.48 ± 0.32* |
| 0.02 | 2.48 ± 0.04 |
| 0.2 | 2.18 ± 0.12 |
| 2 | 2.06 ± 0.04 |
| 20 | 1** |

**Table S3: Error rates and fold changes by mutation context for targeted panel sequencing**. Duplex sequencing error rates broken down by mutation context for four healthy donor cfDNA samples, three cancer patient cfDNA samples, and five FFPE tumor biopsies. The samples were treated with Duplex-Repair and conventional ER/AT.

| Specimen | Patient ID | Mutation Context | Error Rate  Conventional ER/AT | Error Rate  Duplex-  Repair | Error Rate  Fold Decrease | CI (95%) |
| --- | --- | --- | --- | --- | --- | --- |
| cfDNA | HD_67 | T>G | 3.83E-08 | 4.38E-08 | -1.14 | [-2.06, -0.23] |
| cfDNA | HD_67 | T>C | 9.70E-08 | 1.38E-07 | -1.42 | [-2.29, -0.55] |
| cfDNA | HD_67 | T>A | 3.83E-08 | 2.41E-08 | 1.59 | [0.89, 2.29] |
| cfDNA | HD_67 | C>A | 2.56E-07 | 5.71E-08 | 4.47 | [3.98, 4.97] |
| cfDNA | HD_67 | C>G | 6.24E-08 | 7.79E-08 | -1.25 | [-2.13, -0.36] |
| cfDNA | HD_67 | C>T | 3.24E-07 | 3.43E-07 | -1.06 | [-2.04, -0.07] |
| cfDNA | HD_77 | T>G | 4.87E-08 | 3.42E-08 | 1.42 | [0.63, 2.22] |
| cfDNA | HD_77 | T>C | 1.46E-07 | 1.48E-07 | -1.01 | [-2.01, -0.02] |
| cfDNA | HD_77 | T>A | 4.87E-08 | 4.11E-08 | 1.19 | [0.28, 2.09] |
| cfDNA | HD_77 | C>A | 1.10E-07 | 4.85E-08 | 2.26 | [1.61, 2.91] |
| cfDNA | HD_77 | C>G | 6.45E-08 | 3.57E-08 | 1.81 | [1.12, 2.49] |
| cfDNA | HD_77 | C>T | 1.18E-06 | 2.55E-07 | 4.65 | [3.93, 5.37] |
| cfDNA | HD_78 | T>G | 1.80E-08 | 1.93E-08 | -1.07 | [-2.01, -0.13] |
| cfDNA | HD_78 | T>C | 4.61E-08 | 3.30E-08 | 1.39 | [0.60, 2.19] |
| cfDNA | HD_78 | T>A | 2.40E-08 | 2.20E-08 | 1.09 | [0.17, 2.02] |
| cfDNA | HD_78 | C>A | 1.39E-07 | 1.94E-08 | 7.19 | [7.00, 7.38] |
| cfDNA | HD_78 | C>G | 3.98E-08 | 1.94E-08 | 2.05 | [1.54, 2.57] |
| cfDNA | HD_78 | C>T | 4.03E-07 | 7.43E-08 | 5.42 | [4.94, 5.90] |
| cfDNA | HD_79 | T>G | 1.10E-07 | 1.15E-08 | 9.5 | [9.44, 9.55] |
| cfDNA | HD_79 | T>C | 1.34E-07 | 4.79E-08 | 2.8 | [2.34, 3.26] |
| cfDNA | HD_79 | T>A | 7.46E-08 | 4.61E-08 | 1.62 | [0.98, 2.26] |
| cfDNA | HD_79 | C>A | 2.39E-07 | 1.31E-07 | 1.82 | [1.09, 2.54] |
| cfDNA | HD_79 | C>G | 1.42E-07 | 5.15E-08 | 2.75 | [2.32, 3.18] |
| cfDNA | HD_79 | C>T | 1.06E-06 | 5.13E-07 | 2.07 | [1.25, 2.90] |
| cfDNA | 05055_33 | C>A | 1.07E-07 | 2.20E-07 | -2.06 | [-2.42, -1.71] |
| cfDNA | 05055_33 | C>G | 1.07E-07 | 7.34E-08 | 1.45 | [1.00, 1.90] |
| cfDNA | 05055_33 | C>T | 2.45E-06 | 4.40E-07 | 5.57 | [5.32, 5.81] |
| cfDNA | 05055_33 | T>A | 3.90E-08 | 6.92E-08 | -1.77 | [-2.03, -1.52] |
| cfDNA | 05055_33 | T>C | 7.81E-08 | 2.08E-07 | -2.66 | [-2.89, -2.43] |
| cfDNA | 05055_33 | T>G | 3.90E-08 | 6.92E-08 | -1.77 | [-2.03, -1.52] |
| cfDNA | 05055_48 | C>A | 5.25E-07 | 0.00E+00 | inf | [0.00, 0.00] |
| cfDNA | 05055_48 | C>G | 2.02E-08 | 1.06E-07 | -5.25 | [-5.28, -5.23] |
| cfDNA | 05055_48 | C>T | 7.89E-06 | 6.01E-07 | 13.13 | [12.85, 13.42] |
| cfDNA | 05055_48 | T>A | 1.60E-08 | 5.74E-08 | -3.59 | [-3.64, -3.54] |
| cfDNA | 05055_48 | T>C | 7.99E-08 | 1.15E-07 | -1.44 | [-2.06, -0.82] |
| cfDNA | 05055_48 | T>G | 3.20E-08 | 2.87E-08 | 1.11 | [0.34, 1.89] |
| cfDNA | 05055_73 | C>A | 4.39E-07 | 1.17E-07 | 3.77 | [3.57, 3.96] |
| cfDNA | 05055_73 | C>G | 2.31E-08 | 0.00E+00 | inf | [0.00, 0.00] |
| cfDNA | 05055_73 | C>T | 2.54E-06 | 5.83E-07 | 4.36 | [3.91, 4.81] |
| cfDNA | 05055_73 | T>A | 3.85E-08 | 3.32E-08 | 1.16 | [0.46, 1.86] |
| cfDNA | 05055_73 | T>C | 3.85E-08 | 3.32E-08 | 1.16 | [0.46, 1.86] |
| cfDNA | 05055_73 | T>G | 1.92E-08 | 0.00E+00 | inf | [0.00, 0.00] |
| FFPE Biopsy | 05055_106 | C>A | 3.80E-06 | 2.59E-07 | 14.68 | [14.41, 14.95] |
| FFPE Biopsy | 05055_106 | C>G | 1.43E-06 | 2.59E-07 | 5.54 | [5.14, 5.94] |
| FFPE Biopsy | 05055_106 | C>T | 5.33E-05 | 1.98E-05 | 2.7 | [1.77, 3.63] |
| FFPE Biopsy | 05055_106 | T>A | 5.48E-07 | 3.35E-07 | 1.64 | [0.87, 2.40] |
| FFPE Biopsy | 05055_106 | T>C | 1.24E-06 | 1.41E-06 | -1.14 | [-2.10, -0.17] |
| FFPE Biopsy | 05055_106 | T>G | 1.69E-07 | 0.00E+00 | inf | [0.00, 0.00] |
| FFPE Biopsy | 05055_129 | C>A | 4.36E-06 | 2.07E-07 | 21.09 | [20.87, 21.30] |
| FFPE Biopsy | 05055_129 | C>G | 2.13E-06 | 2.84E-07 | 7.47 | [7.07, 7.88] |
| FFPE Biopsy | 05055_129 | C>T | 5.17E-05 | 1.25E-05 | 4.14 | [3.24, 5.05] |
| FFPE Biopsy | 05055_129 | T>A | 3.25E-07 | 9.53E-08 | 3.41 | [3.03, 3.79] |
| FFPE Biopsy | 05055_129 | T>C | 1.11E-06 | 7.87E-07 | 1.41 | [0.52, 2.31] |
| FFPE Biopsy | 05055_129 | T>G | 1.37E-07 | 2.38E-08 | 5.77 | [5.70, 5.84] |
| FFPE Biopsy | 05055_2 | C>A | 1.16E-05 | 6.10E-07 | 19.02 | [18.72, 19.31] |
| FFPE Biopsy | 05055_2 | C>G | 6.14E-06 | 9.41E-07 | 6.53 | [6.01, 7.04] |
| FFPE Biopsy | 05055_2 | C>T | 1.89E-04 | 2.53E-05 | 7.5 | [6.63, 8.38] |
| FFPE Biopsy | 05055_2 | T>A | 9.43E-07 | 7.24E-07 | 1.3 | [0.44, 2.17] |
| FFPE Biopsy | 05055_2 | T>C | 4.91E-06 | 1.96E-06 | 2.5 | [1.74, 3.26] |
| FFPE Biopsy | 05055_2 | T>G | 1.17E-06 | 1.03E-07 | 11.33 | [11.27, 11.39] |
| FFPE Biopsy | 05055_73 | C>A | 4.47E-06 | 2.13E-07 | 21.01 | [20.81, 21.20] |
| FFPE Biopsy | 05055_73 | C>G | 1.71E-06 | 2.74E-07 | 6.24 | [5.84, 6.65] |
| FFPE Biopsy | 05055_73 | C>T | 3.71E-05 | 9.63E-06 | 3.85 | [2.96, 4.74] |
| FFPE Biopsy | 05055_73 | T>A | 2.67E-07 | 1.86E-07 | 1.43 | [0.65, 2.22] |
| FFPE Biopsy | 05055_73 | T>C | 1.08E-06 | 7.32E-07 | 1.47 | [0.60, 2.35] |
| FFPE Biopsy | 05055_73 | T>G | 1.78E-07 | 0.00E+00 | inf | [0.00, 0.00] |
| FFPE Biopsy | 05055_95 | C>A | 6.60E-06 | 6.79E-07 | 9.73 | [9.31, 10.15] |
| FFPE Biopsy | 05055_95 | C>G | 2.91E-06 | 9.28E-07 | 3.14 | [2.48, 3.80] |
| FFPE Biopsy | 05055_95 | C>T | 1.12E-04 | 2.77E-05 | 4.05 | [3.14, 4.97] |
| FFPE Biopsy | 05055_95 | T>A | 4.67E-07 | 3.69E-07 | 1.27 | [0.41, 2.12] |
| FFPE Biopsy | 05055_95 | T>C | 2.42E-06 | 2.28E-06 | 1.06 | [0.08, 2.05] |
| FFPE Biopsy | 05055_95 | T>G | 5.77E-07 | 1.52E-07 | 3.8 | [3.48, 4.12] |

**Table S4: Sequencing metrics for all samples profiled by targeted panel sequencing**.

| Specimen | Patient ID | DNA damage inducers | ER/AT method | Number of raw reads | On target rates | Number of duplex bases evaluated | Number of base errors | Error Rate | CI (95%) |
| --- | --- | --- | --- | --- | --- | --- | --- | --- | --- |
| cfDNA | HD_78 | 0uM_CuCl2/H2O2+0mU_DNase1 | Conv. ER/AT | 1.29E+08 | 0.98172 | 12216528 | 1 | 8.19E-08 | [1.44e-08, 4.64e-07] |
| cfDNA | HD_78 | 0uM_CuCl2/H2O2+0.2mU_DNase1 | Conv. ER/AT | 1.54E+08 | 0.982728 | 10417986 | 3 | 2.88E-07 | [9.79e-08, 8.47e-07] |
| cfDNA | HD_78 | 0uM_CuCl2/H2O2+2mU_DNase1 | Conv. ER/AT | 1.39E+08 | 0.983843 | 9406423 | 5 | 5.32E-07 | [2.27e-07, 1.24e-06] |
| cfDNA | HD_78 | 2uM_CuCl2/H2O2+0mU_DNase1 | Conv. ER/AT | 1.58E+08 | 0.982447 | 10542247 | 4 | 3.79E-07 | [1.48e-07, 9.76e-07] |
| cfDNA | HD_78 | 2uM_CuCl2/H2O2+0.2mU_DNase1 | Conv. ER/AT | 1.36E+08 | 0.982074 | 11439892 | 5 | 4.37E-07 | [1.87e-07, 1.02e-06] |
| cfDNA | HD_78 | 2uM_CuCl2/H2O2+2mU_DNase1 | Conv. ER/AT | 1.85E+08 | 0.983726 | 7559620 | 5 | 6.61E-07 | [2.83e-07, 1.55e-06] |
| cfDNA | HD_78 | 100uM_CuCl2/H2O2+0mU_DNase1 | Conv. ER/AT | 1.33E+08 | 0.982054 | 11668760 | 5 | 4.28E-07 | [1.83e-07, 1.00e-06] |
| cfDNA | HD_78 | 100uM_CuCl2/H2O2+0.2mU_DNase1 | Conv. ER/AT | 1.47E+08 | 0.981823 | 10624035 | 7 | 6.59E-07 | [3.19e-07, 1.36e-06] |
| cfDNA | HD_78 | 100uM_CuCl2/H2O2+2mU_DNase1 | Conv. ER/AT | 1.66E+08 | 0.983928 | 8380284 | 7 | 8.35E-07 | [4.05e-07, 1.72e-06] |
| cfDNA | HD_78 | 0uM_CuCl2/H2O2+0mU_DNase1 | Conv. ER/AT | 1.51E+08 | 0.981821 | 13171944 | 3 | 2.28E-07 | [7.75e-08, 6.70e-07] |
| cfDNA | HD_78 | 0uM_CuCl2/H2O2+0.2mU_DNase1 | Conv. ER/AT | 1.30E+08 | 0.982407 | 11141949 | 3 | 2.69E-07 | [9.16e-08, 7.92e-07] |
| cfDNA | HD_78 | 0uM_CuCl2/H2O2+2mU_DNase1 | Conv. ER/AT | 1.66E+08 | 0.983527 | 10042644 | 6 | 5.97E-07 | [2.74e-07, 1.30e-06] |
| cfDNA | HD_78 | 2uM_CuCl2/H2O2+0mU_DNase1 | Conv. ER/AT | 1.20E+08 | 0.982558 | 10956968 | 1 | 9.13E-08 | [1.61e-08, 5.17e-07] |
| cfDNA | HD_78 | 2uM_CuCl2/H2O2+0.2mU_DNase1 | Conv. ER/AT | 1.67E+08 | 0.982041 | 10056176 | 2 | 1.99E-07 | [5.45e-08, 7.25e-07] |
| cfDNA | HD_78 | 2uM_CuCl2/H2O2+2mU_DNase1 | Conv. ER/AT | 1.95E+08 | 0.983867 | 8965981 | 3 | 3.35E-07 | [1.14e-07, 9.84e-07] |
| cfDNA | HD_78 | 100uM_CuCl2/H2O2+0mU_DNase1 | Conv. ER/AT | 1.30E+08 | 0.982483 | 12904331 | 3 | 2.32E-07 | [7.91e-08, 6.84e-07] |
| cfDNA | HD_78 | 100uM_CuCl2/H2O2+0.2mU_DNase1 | Conv. ER/AT | 1.30E+08 | 0.981903 | 10159501 | 6 | 5.91E-07 | [2.71e-07, 1.29e-06] |
| cfDNA | HD_78 | 100uM_CuCl2/H2O2+2mU_DNase1 | Conv. ER/AT | 1.52E+08 | 0.984147 | 6953914 | 7 | 1.01E-06 | [4.88e-07, 2.08e-06] |
| cfDNA | HD_78 | 0uM_CuCl2/H2O2+0mU_DNase1 | Conv. ER/AT | 1.49E+08 | 0.980883 | 12742674 | 4 | 3.14E-07 | [1.22e-07, 8.07e-07] |
| cfDNA | HD_78 | 0uM_CuCl2/H2O2+0.2mU_DNase1 | Conv. ER/AT | 1.39E+08 | 0.981531 | 12832585 | 3 | 2.34E-07 | [7.95e-08, 6.87e-07] |
| cfDNA | HD_78 | 0uM_CuCl2/H2O2+2mU_DNase1 | Conv. ER/AT | 1.48E+08 | 0.983953 | 6998068 | 2 | 2.86E-07 | [7.84e-08, 1.04e-06] |
| cfDNA | HD_78 | 2uM_CuCl2/H2O2+0mU_DNase1 | Conv. ER/AT | 1.53E+08 | 0.981475 | 13027274 | 0 | 0 | [0.00e+00, 2.95e-07] |
| cfDNA | HD_78 | 2uM_CuCl2/H2O2+0.2mU_DNase1 | Conv. ER/AT | 1.54E+08 | 0.982044 | 11278606 | 1 | 8.87E-08 | [1.57e-08, 5.02e-07] |
| cfDNA | HD_78 | 2uM_CuCl2/H2O2+2mU_DNase1 | Conv. ER/AT | 1.82E+08 | 0.982618 | 8682012 | 5 | 5.76E-07 | [2.46e-07, 1.35e-06] |
| cfDNA | HD_78 | 100uM_CuCl2/H2O2+0mU_DNase1 | Conv. ER/AT | 1.37E+08 | 0.981088 | 11267015 | 5 | 4.44E-07 | [1.90e-07, 1.04e-06] |
| cfDNA | HD_78 | 100uM_CuCl2/H2O2+0.2mU_DNase1 | Conv. ER/AT | 1.33E+08 | 0.980935 | 8874034 | 3 | 3.38E-07 | [1.15e-07, 9.94e-07] |
| cfDNA | HD_78 | 100uM_CuCl2/H2O2+2mU_DNase1 | Conv. ER/AT | 1.27E+08 | 0.983378 | 7319479 | 5 | 6.83E-07 | [2.92e-07, 1.60e-06] |
| cfDNA | damaged_HD_78_cfDNA | 100uM_CuCl2/H2O2_+_2mU_DNase1 | Duplex-Repair | 5.34E+07 | 0.988432 | 44538494 | 13 | 2.92E-07 | [1.71e-07, 4.99e-07] |
| cfDNA | damaged_HD_78_cfDNA | 100uM_CuCl2/H2O2_+_2mU_DNase2 | Duplex-Repair | 6.42E+07 | 0.988659 | 43911733 | 21 | 4.78E-07 | [3.13e-07, 7.31e-07] |
| cfDNA | damaged_HD_78_cfDNA | 100uM_CuCl2/H2O2_+_2mU_DNase3 | Duplex-Repair | 6.92E+07 | 0.988647 | 50237066 | 18 | 3.58E-07 | [2.27e-07, 5.66e-07] |
| cfDNA | HD_78 | NA | Duplex-Repair | 1.12E+08 | 0.983891 | 220967435 | 20 | 9.05E-08 | [5.86e-08, 1.40e-07] |
| cfDNA | HD_78 | NA | Duplex-Repair | 1.22E+08 | 0.982716 | 205270556 | 24 | 1.17E-07 | [7.86e-08, 1.74e-07] |
| cfDNA | HD_78 | NA | Duplex-Repair | 1.15E+08 | 0.983438 | 246649702 | 24 | 9.73E-08 | [6.54e-08, 1.45e-07] |
| cfDNA | 05055_33 | NA | Duplex-Repair | 1.48E+07 | 0.965585 | 28090417 | 15 | 5.34E-07 | [3.24e-07, 8.81e-07] |
| cfDNA | 05055_33 | NA | Conv. ER/AT | 3.96E+07 | 0.970814 | 98148695 | 133 | 1.36E-06 | [1.14e-06, 1.61e-06] |
| cfDNA | 05055_48 | NA | Duplex-Repair | 3.91E+07 | 0.986819 | 63145388 | 27 | 4.28E-07 | [2.94e-07, 6.22e-07] |
| cfDNA | 05055_48 | NA | Conv. ER/AT | 6.12E+07 | 0.980437 | 112122125 | 426 | 3.80E-06 | [3.46e-06, 4.18e-06] |
| cfDNA | 05055_73 | NA | Duplex-Repair | 3.39E+07 | 0.983176 | 55844344 | 20 | 3.58E-07 | [2.32e-07, 5.53e-07] |
| cfDNA | 05055_73 | NA | Conv. ER/AT | 3.46E+07 | 0.981357 | 95264860 | 135 | 1.42E-06 | [1.20e-06, 1.68e-06] |
| FFPE Tumor Biopsy | 05055_106 | NA | Conv. ER/AT | 5.36E+07 | 0.98884 | 20006860 | 521 | 2.60E-05 | [2.39e-05, 2.84e-05] |
| FFPE Tumor Biopsy | 05055_106 | NA | Conv. ER/AT | 6.61E+07 | 0.988419 | 23707349 | 665 | 2.81E-05 | [2.60e-05, 3.03e-05] |
| FFPE Tumor Biopsy | 05055_106 | NA | Conv. ER/AT | 9.51E+07 | 0.988755 | 43482673 | 1235 | 2.84E-05 | [2.69e-05, 3.00e-05] |
| FFPE Tumor Biopsy | 05055_106 | NA | Duplex-Repair | 6.73E+07 | 0.991287 | 49319461 | 467 | 9.47E-06 | [8.65e-06, 1.04e-05] |
| FFPE Tumor Biopsy | 05055_106 | NA | Duplex-Repair | 8.13E+07 | 0.991346 | 53118736 | 538 | 1.01E-05 | [9.31e-06, 1.10e-05] |
| FFPE Tumor Biopsy | 05055_106 | NA | Duplex-Repair | 6.85E+07 | 0.991322 | 50713014 | 551 | 1.09E-05 | [9.99e-06, 1.18e-05] |
| FFPE Tumor Biopsy | 05055_95 | NA | Conv. ER/AT | 8.51E+07 | 0.990015 | 17146216 | 879 | 5.13E-05 | [4.80e-05, 5.48e-05] |
| FFPE Tumor Biopsy | 05055_95 | NA | Conv. ER/AT | 7.61E+07 | 0.990807 | 24648513 | 1564 | 6.35E-05 | [6.04e-05, 6.67e-05] |
| FFPE Tumor Biopsy | 05055_95 | NA | Conv. ER/AT | 8.33E+07 | 0.990089 | 27917450 | 1734 | 6.21E-05 | [5.93e-05, 6.51e-05] |
| FFPE Tumor Biopsy | 05055_95 | NA | Duplex-Repair | 8.51E+07 | 0.991906 | 32413963 | 530 | 1.64E-05 | [1.50e-05, 1.78e-05] |
| FFPE Tumor Biopsy | 05055_95 | NA | Duplex-Repair | 7.40E+07 | 0.991789 | 28687673 | 471 | 1.64E-05 | [1.50e-05, 1.80e-05] |
| FFPE Tumor Biopsy | 05055_95 | NA | Duplex-Repair | 7.44E+07 | 0.991882 | 29219082 | 421 | 1.44E-05 | [1.31e-05, 1.59e-05] |
| FFPE Tumor Biopsy | 05055_2 | NA | Conv. ER/AT | 6.47E+07 | 0.984017 | 13700238 | 1078 | 7.87E-05 | [7.41e-05, 8.35e-05] |
| FFPE Tumor Biopsy | 05055_2 | NA | Conv. ER/AT | 5.20E+07 | 0.985261 | 23873009 | 2729 | 0.00011431 | [1.10e-04, 1.19e-04] |
| FFPE Tumor Biopsy | 05055_2 | NA | Conv. ER/AT | 7.87E+07 | 0.986823 | 24256287 | 2853 | 0.00011762 | [1.13e-04, 1.22e-04] |
| FFPE Tumor Biopsy | 05055_2 | NA | Duplex-Repair | 1.00E+08 | 0.992637 | 26565686 | 469 | 1.77E-05 | [1.61e-05, 1.93e-05] |
| FFPE Tumor Biopsy | 05055_2 | NA | Duplex-Repair | 1.03E+08 | 0.992481 | 27540728 | 457 | 1.66E-05 | [1.51e-05, 1.82e-05] |
| FFPE Tumor Biopsy | 05055_2 | NA | Duplex-Repair | 9.27E+07 | 0.992096 | 14224743 | 209 | 1.47E-05 | [1.28e-05, 1.68e-05] |
| FFPE Tumor Biopsy | 05055_73 | NA | Conv. ER/AT | 9.77E+07 | 0.988559 | 49482848 | 988 | 2.00E-05 | [1.88e-05, 2.13e-05] |
| FFPE Tumor Biopsy | 05055_73 | NA | Conv. ER/AT | 8.96E+07 | 0.988492 | 45128942 | 860 | 1.91E-05 | [1.78e-05, 2.04e-05] |
| FFPE Tumor Biopsy | 05055_73 | NA | Conv. ER/AT | 8.27E+07 | 0.991062 | 51361631 | 1184 | 2.31E-05 | [2.18e-05, 2.44e-05] |
| FFPE Tumor Biopsy | 05055_73 | NA | Duplex-Repair | 9.48E+07 | 0.991359 | 46828201 | 251 | 5.36E-06 | [4.74e-06, 6.07e-06] |
| FFPE Tumor Biopsy | 05055_73 | NA | Duplex-Repair | 9.98E+07 | 0.992118 | 43268486 | 197 | 4.55E-06 | [3.96e-06, 5.23e-06] |
| FFPE Tumor Biopsy | 05055_73 | NA | Duplex-Repair | 1.03E+08 | 0.991284 | 50748145 | 286 | 5.64E-06 | [5.02e-06, 6.33e-06] |
| FFPE Tumor Biopsy | 05055_129 | NA | Conv. ER/AT | 6.19E+07 | 0.990651 | 38198285 | 790 | 2.07E-05 | [1.93e-05, 2.22e-05] |
| FFPE Tumor Biopsy | 05055_129 | NA | Conv. ER/AT | 5.70E+07 | 0.988617 | 46259759 | 1411 | 3.05E-05 | [2.90e-05, 3.21e-05] |
| FFPE Tumor Biopsy | 05055_129 | NA | Conv. ER/AT | 8.37E+07 | 0.98878 | 67552550 | 2111 | 3.12E-05 | [2.99e-05, 3.26e-05] |
| FFPE Tumor Biopsy | 05055_129 | NA | Duplex-Repair | 6.32E+07 | 0.990994 | 46410054 | 300 | 6.46E-06 | [5.77e-06, 7.24e-06] |
| FFPE Tumor Biopsy | 05055_129 | NA | Duplex-Repair | 6.47E+07 | 0.990846 | 47452054 | 315 | 6.64E-06 | [5.94e-06, 7.41e-06] |
| FFPE Tumor Biopsy | 05055_129 | NA | Duplex-Repair | 6.50E+07 | 0.991667 | 67397721 | 463 | 6.87E-06 | [6.27e-06, 7.52e-06] |
| cfDNA | HD_67 | NA | Conv. ER/AT | 1.15E+08 | 0.987253 | 287241318 | 94 | 3.27E-07 | [2.67e-07, 4.00e-07] |
| cfDNA | HD_67 | NA | Conv. ER/AT | 1.21E+08 | 0.987747 | 228373435 | 100 | 4.38E-07 | [3.60e-07, 5.33e-07] |
| cfDNA | HD_67 | NA | Conv. ER/AT | 1.07E+08 | 0.987641 | 212510412 | 90 | 4.24E-07 | [3.45e-07, 5.21e-07] |
| cfDNA | HD_67 | NA | Duplex-Repair | 155620562 | 0.980312 | 326796016 | 95 | 2.91E-07 | [2.38e-07, 3.55e-07] |
| cfDNA | HD_67 | NA | Duplex-Repair | 113886380 | 0.980136 | 242483444 | 102 | 4.21E-07 | [3.47e-07, 5.11e-07] |
| cfDNA | HD_67 | NA | Duplex-Repair | 127505618 | 0.980876 | 272308087 | 81 | 2.97E-07 | [2.39e-07, 3.70e-07] |
| cfDNA | HD_77 | NA | Conv. ER/AT | 159711982 | 0.987585 | 459509299 | 318 | 6.92E-07 | [6.20e-07, 7.72e-07] |
| cfDNA | HD_77 | NA | Conv. ER/AT | 134944370 | 0.988378 | 247081170 | 201 | 8.13E-07 | [7.09e-07, 9.34e-07] |
| cfDNA | HD_77 | NA | Conv. ER/AT | 126086546 | 0.987581 | 271681550 | 238 | 8.76E-07 | [7.72e-07, 9.95e-07] |
| cfDNA | HD_77 | NA | Duplex-Repair | 168105150 | 0.976197 | 320539624 | 96 | 2.99E-07 | [2.45e-07, 3.66e-07] |
| cfDNA | HD_77 | NA | Duplex-Repair | 147197612 | 0.97676 | 287680401 | 74 | 2.57E-07 | [2.05e-07, 3.23e-07] |
| cfDNA | HD_77 | NA | Duplex-Repair | 113851702 | 0.973208 | 222307450 | 61 | 2.74E-07 | [2.14e-07, 3.52e-07] |
| cfDNA | HD_79 | NA | Conv. ER/AT | 83749956 | 0.978007 | 20839707 | 26 | 1.25E-06 | [8.51e-07, 1.83e-06] |
| cfDNA | HD_79 | NA | Conv. ER/AT | 89172830 | 0.977934 | 22898460 | 25 | 1.09E-06 | [7.40e-07, 1.61e-06] |
| cfDNA | HD_79 | NA | Conv. ER/AT | 146134132 | 0.983414 | 152767879 | 109 | 7.14E-07 | [5.92e-07, 8.61e-07] |
| cfDNA | HD_79 | NA | Duplex-Repair | 178781140 | 0.977662 | 251559928 | 88 | 3.50E-07 | [2.84e-07, 4.31e-07] |
| cfDNA | HD_79 | NA | Duplex-Repair | 90429248 | 0.974458 | 22015921 | 35 | 1.59E-06 | [1.14e-06, 2.21e-06] |
| cfDNA | HD_79 | NA | Duplex-Repair | 160665758 | 0.977917 | 241459115 | 74 | 3.06E-07 | [2.44e-07, 3.85e-07] |
